# Volumetric reconstruction of main *Caenorhabditis elegans* neuropil at two different time points

**DOI:** 10.1101/485771

**Authors:** Christopher A. Brittin, Steven J. Cook, David H. Hall, Scott W. Emmons, Netta Cohen

**Author notes:** Senior authors (Scott W. Emmons, Netta Cohen).

## Abstract

Detailed knowledge of both synaptic connectivity and the spatial proximity of neurons is crucial for understanding wiring specificity in the nervous system. Here, we volumetrically reconstructed the *C. elegans* nerve ring from legacy serial-sectioned electron micrographs at two distinct time points: the L4 and young adult. The new volumetric reconstructions provide detailed spatial and morphological information of neural processes in the nerve ring. Our analysis suggests that the nerve ring exhibits three levels of wiring specificity: spatial, synaptic and subcellular. Neuron classes innervate well defined neighborhoods and aggregate functionally similar synapses to support distinct computational pathways. Connectivity fractions vary based on neuron class and synapse type. We find that the variability in process placement accounts for less than 20% of the variability in synaptic connectivity and models based only on spatial information cannot account for the reproducibility of synaptic connections among homologous neurons. This suggests that additional, non-spatial factors also contribute to synaptic and subcellular specificity. With this in mind, we conjecture that a spatially constrained, genetic model could provide sufficient synaptic specificity. Using a model of cell-specific combinatorial genetic expression, we show that additional specificity, such as sub-cellular domains or alternative splicing, would be required to reproduce the wiring specificity in the nerve ring.

## Introduction

Wiring specificity, the stereotypic patterns of synaptic connectivity in neural circuits, is a common feature of nervous systems across species (Sanes & Zipursky, 2010). In order to achieve wiring specificity, a nervous system must coordinate both spatial and synaptic specificity. Spatial specificity refers to the anatomical organization of neurons whereby potential synaptic partners are placed in close spatial proximity. The vertebrate neocortex, olfactory bulb and visual system all exhibit a stereotyped multi-layered structure (Baier, 2013; Gilmore & Herrup, 1997; Nagayama et al., 2014; Sanes & Zipursky, 2010) that helps aggregate synapses with similar functional properties to restricted anatomical regions. Synaptic specificity refers to the selectivity of neurons when choosing synaptic partners from the possibly large number of physically adjacent cells. Ultrastructural studies have shown that neurons make stereotypic synaptic partnerships (White et al., 1986; Meinertzhagen & O’Neil, 1991) while displaying relatively low connectivity fractions with neighboring cells (less than 0.25) (Hamos et al., 1987; Escobar et al., 2008; Mishchenko et al., 2010). This suggests that both synaptic and spatial specificity are required, but what are the relative contributions of each to the overall wiring specificity of the nervous system?

If spatial specificity is the major contributor to wiring specificity, then wiring specificity can be modeled statistically once the spatial organization of the nervous system is known. Early models proposed that synaptic patterns could be described by a statistical model(Braitenberg & Schüz, 1998) where the amount of synaptic connectivity is proportional to the amount of contact between axons and dendrites, i.e. Peters’ rule (Rees et al., 2017). There is both experimental and theoretical evidence which suggest that the number of synaptic contacts is indeed proportional to the amount of axo-dendritic contact (Czajkowski et al., 2013; Packer et al., 2013; van Pelt & van Ooyen, 2013). Additional studies also suggest that at least some structural and functional characteristics of connectivity can be inferred from the spatial proximity of neurons (Hill et al., 2012; Reimann et al., 2015). However, other studies suggest that simple statistical models cannot capture the variation in connectivity among different neurons (Mishchenko et al., 2010; Kasthuri et al., 2015; Druckmann et al., 2014). While informative, all of these studies have focused on isolated components of extremely complex neural circuits, which makes it difficult to infer how stereotyped connectivity emerges at the level of a complete neural circuit.

If spatial and synaptic specificity are equally important, then models of wiring specificity will also require detailed knowledge of the molecular events necessary for synapse formation (de Wit & Ghosh, 2015). Classic work by Langley (Langley, 1895) and Sperry (Sperry, 1963) has lead to the “chemoaffinity” hypothesis which states that neurons possess unique cytochemical labels that allow neurons to selectively navigate to their target cells (Meyer, 1998). In order to regulate wiring specificity and synaptic diversity, such surface labels would need to be expressed in unique combinations among distinct neuronal populations (de Wit & Ghosh, 2015). It has been postulated that the molecular diversification of cell adhesion molecules (CAMs) could provide such combinatoric expression (Zipursky & Sanes, 2010; de Wit & Ghosh, 2015). However, to our knowledge, a combinatorial expression model has yet to be formalized in a way that can be tested against a biological neural circuit of known synaptic connectivity at single synapse resolution.

The nematode *C. elegans* offers unique advantages for studying wiring specificity. The worm has a small well-defined nervous system with just 302 neurons in the adult hermaphrodite and neuron classes can be identified based on morphology and cell placement (Cook et al., 2018; Varshney et al., 2011; White et al., 1986). *C. elegans* neurons have simple morphologies and only make *en passant* synapses due to lack of axon terminals. Moreover, the bilateral symmetry of the worm can be exploited in order to asses the variability in neuronal placement and connectivity. However, while the *C. elegans* wiring diagram has been known for over 30 years (White et al., 1986), the spatial proximity of neurons has only been partially characterized based on subsampled data taken from larval stage 4 (L4) electron micrographs (White et al., 1983; Durbin, 1987). Because the data is subsampled, contact between adjacent cells is underestimated. Because the data is taken from the L4, comparison with adult synaptic connectivity, the primary source of the *C. elegans* canonical wiring diagram (Varshney et al., 2011; Cook et al., 2018), is difficult. To remedy this, we have used legacy serial section electron micrographs (White et al., 1986) to volumetrically reconstruct a L4 and an adult nerve ring. The nerve ring is a cycloneuralian brain (Richter et al., 2010), a neuropile of uniform thickness that surrounds the pharynx of the worm. We find that the *C. elegans* nerve ring is organized into a quasi-layered structure that separates distinct computational pathways by aggregating functionally similar synapses. We show that neuron classes innervate distinct neighborhoods, suggesting that process placement is specified. However, using the recently updated synaptic wiring diagram (Cook et al., 2018), we show that the spatial specificity is not sufficient to account for the reproducibility of synaptic connectivity, suggesting that non-spatial factors also contribute to wiring specificity. Finally, we test three variations of a combinatorial CAM expression model and show that under certain conditions such a model could account for wiring specificity in the nerve ring. Collectively, these results suggest that both spatial and synaptic specificity are critical for overall wiring specificity in the *C. elegans* nerve ring.

## Results

### Volumetric reconstruction of C. *elegans* nerve ring

We volumetrically reconstructed the nerve rings of two animals using previously published serial section electron micrographs (EMs) (White et al., 1986), one from the young adult and one from a larval stage 4 (L4) animal (Table S1). Both EM series consist of ~ 90 nm thick sections that approximately span the same 36 *μ*m long volume, starting and ending in the anterior and ventral ganglia, respectively (Figure 1(a)). These correspond to 400 EMs in the L4 animal. In the adult series, all EM sections were analyzed starting in the anterior ganglion but only every other EM section was imaged starting in the ventral ganglion (posterior to the nerve ring), such that only 300 sections were included in the adult data set (see Methods). We corrected for the reduced number of EM sections in our analysis (see Methods), but also found that the qualitative results did not depend on the correction.

**Figure 1.**
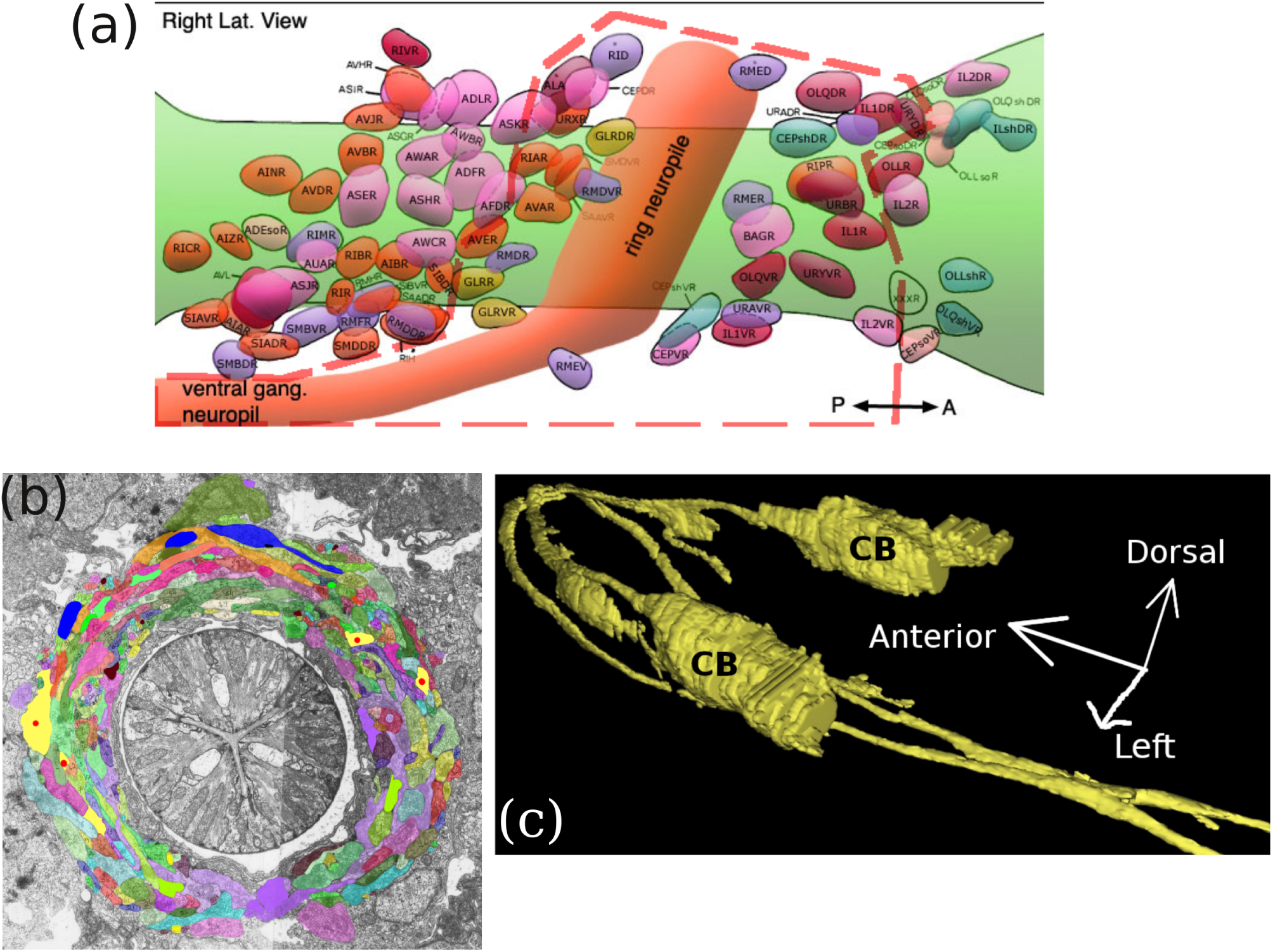
Overview of anatomy and volumetric reconstruction. (a) Nuclei positions of cells that project axons/processes into the nerve ring. All processes projected into the nerve ring were reconstructed. Only cell bodies within the dashed red boundary were reconstructed. (Modified image from wormatlas.org.) (b) A segmented EM taken from the nerve ring. Neurons are manually segmented and each neuron assigned a different color. Segmentation was performed for 300 and 400 EM sections in the adult and L4, respectively. Red dots indicate processes of the AVA neurons. (c) A 3D reconstruction of neurons AVAL and AVAR generated from the segmentation data. CB: cell body.

We volumetrically reconstructed neurons by using TrakEM2 (Cardona et al., 2012) to manually segment processes and somata in each EM section (Figure 1(b)). In both the L4 and the adult, there are 181 neurons from neuron classes that send axons/processes into the nerve ring (Table S1). Most neuron classes consist of 1–3 contralateral (left/right homologous) pairs of neurons that share a similar lineage history (Sulston et al., 1983) and are functionally, genetically and morphologically similar (White et al., 1986; Hobert et al., 2002). All axons/processes in the nerve ring and some somata in the anterior and ventral ganglia were segmented. Dendritic processes extending from the amphid and labial sensory neurons towards the nose were not segmented because these portions have very few synapses and therefore were not of immediate interest.

We developed a web application to view the volumetric reconstructions which is available at wormwiring.org. The app shows that neuron processes exhibit a wide range of complex morphologies in the nerve ring. Neuron morphologies range from cylindrical and tube-like to flat with a wide ribbon-like cross-sectional area (Figures 2(a) and S1). Even the morphology of a single neuron can vary greatly along the length of its process (Figures 1(c) and S1). Some neurons exhibit branching, while many do not. Some neuron processes grow contralaterally across the nerve ring commissure, while many processes only grow ipsilaterally.

**Figure 2.**
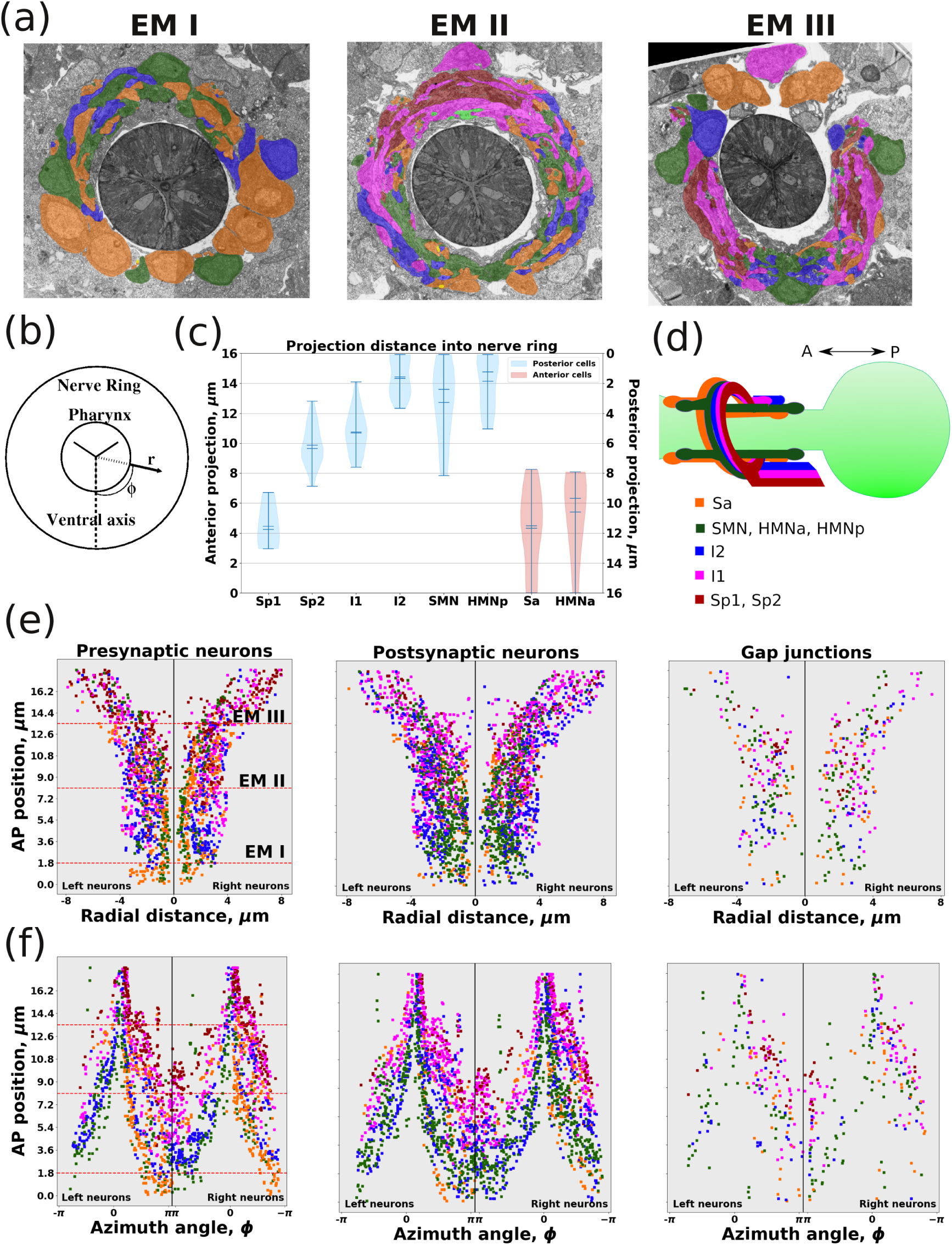
Nerve ring is organized into layered-like structure. (a) Representative EM sections taken from *z* positions given in (e). Neuron colors given in (c). (b) Positions of neurons/synapses are given in terms of cylindrical coordinates (*r*, *ϕ*, *z*). *r* is the distance from the outer edge of pharynx. *ϕ* is azimuth angle with respect to the ventral axis; +*ϕ* is clockwise; −*ϕ* is anti-clockwise. *z* is the position along the anterior-posterior (AP) axis; positive *z* moves in the posterior directions. (c) The projection distance of cells into the nerve ring. Posterior nuclei (blue) project anteriorly. Anterior nuclei (red) project posteriorly. Cells grouped based on function and length of projection (see main text). Vertical bars are the range of projections for given cell group. Width of bar is the fraction of cells at given length. Middle ticks are the mean and median. (d) Illustrations of how projections create layers in the nerve ring. (e) The (*r*, *z*) map of synapses. Colors represent the presynaptic (left), postsynaptic (middle) and gap junction (right) partners. Maps are split to show the left and right side of nerve ring. (f) The (*ϕ*, *z*) map of synapses.

We constructed an algorithm that quantified adjacency directly from the TrakEM2 segmentation. The algorithm identifies all adjacent neurons, identifies where the neurons make physical contact and quantifies the total surface area of contact between them (see Methods). When the cells are properly segmented with TrakEM2, the algorithm outperformed manual identification of physically adjacent cells by experts (see Methods). The cell adjacencies identified by the algorithm are also consistent with previously reported cell adjacencies (White et al., 1983).

### The nerve ring is spatially organized to support distinct computational pathways

We asked how the spatial organization of processes in the nerve ring contributes to the organization of computational pathways. Two structural features stand out from our analysis. First, projections from different anatomical and functional groups of neurons form a layered structure within the nerve ring. Second, mechanosensory and amphid sensory synaptic pathways are physically distinct.

We used cylindrical coordinates (*r*, *ϕ*, *z*) to characterize the spatial structure of the nerve ring (Figure 2(b)). The radius (*r*) is measured as the distance from the outer edges of the pharynx to the neuron or synapse. The azimuth angle (*ϕ*) is measured with respect to the ventral axis, with positive *ϕ* moving in the clockwise direction. The *z* coordinate gives the position along the anterior-posterior (AP) axis of the worm, with *z* = 0 located just anterior to the nerve ring and positive *z* moving in the posterior direction. We analyzed the spatial organization of the nerve ring with respect to each cylindrical dimension.

We assigned each neuron class to one of seven groups based on function and cell body placement (Table S2). The neuron groups presented here are based on our previous classifications (Cook et al., 2018), but have been slightly altered in order emphasize anatomical rather than functional characteristics. The anterior sensory group (Sa) consists of mechanosensory and O_2_/CO_2_ sensing neurons and have cell bodies anterior to the nerve ring. The posterior sensory group (Sp) consists of the amphid sensory neurons and have cell bodies in the lateral ganglion posterior to the nerve ring. The I1 and I2 groups consist of the first- and second-layer interneurons, respectively. The sublateral motor neuron group (SMN) consists of head motor neurons that also send processes posteriorly down the sublateral cords. The HMN group consists of the remaining head motor neurons. The HMN group can further be subdivided into neurons with cell bodies anterior (HMNa) or posterior (HMNp) to the nerve ring.

We find that these neuron groups project different distances into the nerve ring along the *z*-axis creating a layered-like organization. We define the projection distance as the maximum distance that a process grows into the nerve ring before terminating or reversing direction along the *z*-axis. Note that projection distances are lengths and not positions along z. Furthermore, projection distances are reported with the direction of growth, either anteriorly for posterior cells (Sp, I1, I2, SMN and HMNp) or posteriorly for anterior cells (Sa and HMNa) (Figure 2(c)). Based on projection distances, the amphid sensory neurons and the interneurons create a three-layered structure along the *z*-axis (Figure 2(d)). The second-layer interneurons (I2) project the furthest into the nerve ring, followed by first-layer interneurons (I1) and then amphid sensory neurons (Sp). The amphids contain two subgroups (Sp1 and Sp2) with different projection distances. Interestingly, Sp1 (which includes classes AVM and SDQ) has the shortest projection distances while having the largest distance to reach the nerve ring because its cell bodies are closer to the vulva. Interneurons project further than Sp neurons and also have two subgroups (I1 and I2) with different projection distances. Posterior motor neurons (SMN and HMNp) extend roughly the same distances as interneurons. Anterior cells (Sa and HMNa) extend roughly the same posterior distance into the nerve ring. These different projection distances suggest a possible three layered structure along the anterior-posterior axis for neuron groups Sp, I1 and I2 (Figure 2(d)). The Sa and motor neuron groups project the length of the nerve ring and do not create obvious layers along the AP axis. Instead, the Sp, I1 and I2 groups grow around the Sa and motor neuron groups which grow closer to the pharynx (small *r*). Thus, the layered structure is complex, occurring along both the radial and *z*-axes.

We find that mechanosensory and amphid sensory pathways start at physically distinct regions in the nerve ring before converging onto the motor neurons. We measured the cylindrical coordinate of each synapse and noted the group identity of the pre- and postsynaptic neurons. In the (*r*, *z*) plane, synapses aggregate in a radial pattern. Chemical synapses involving either mechanosensory (Sa) neurons or motor neurons (HMN and SMN) mostly aggregate closer to the pharynx (Figure 2(d); green and yellow dots). This places mechanosensory and motor neurons next to head muscle arms which surround the pharynx and may be intended to reduce the processing steps between touch stimuli and head response. Moving anteriorly in the (*ϕ, z*) plane (Figure 2(e)), amphid sensory neurons mostly innervate first-layer and to a lesser extent second-layer interneurons. First-layer interneurons are followed by second-layer interneurons. Second-layer interneurons mix with first-layer interneurons and motor neurons and bridge the connectivity between the two neuron categories. Thus, amphid sensory signals physically travel further to reach muscle output.

## Relative process placement is specified

We next assessed the importance and reproducibility of process placement in the nerve ring through an analysis of neuronal neighborhoods. We define two neurons as neighbors if they are physically adjacent in at least one EM section. The set of neighbors for neuron *i* is the neighborhood of *i* and the size of the neighborhood is measured by the adjacency degree *d_i_* (the number of cells in the neighborhood of *i* excluding *i*). Neurons exhibit a wide range of adjacency degrees which appear to be correlated with their anatomical grouping (Figure 3(a) and Table S2). Distributions of group adjacency degrees are similar between the L4 and the adult. On average, sensory neurons (Sp and Sa) have a lower degree, motor neurons (SMN and HMNp) have a higher degree and interneurons (I1 and I2) have a wide degree range (Figure 3(a)). The motor neuron group HMNa, consisting of the RME and URA classes, are the exception to this trend and have degree distributions which are similar to the sensory neurons. The HMNa neurons have some of the shortest processes in the nerve ring and mostly innervate the inner radial segments closest to the pharynx.

**Figure 3.**
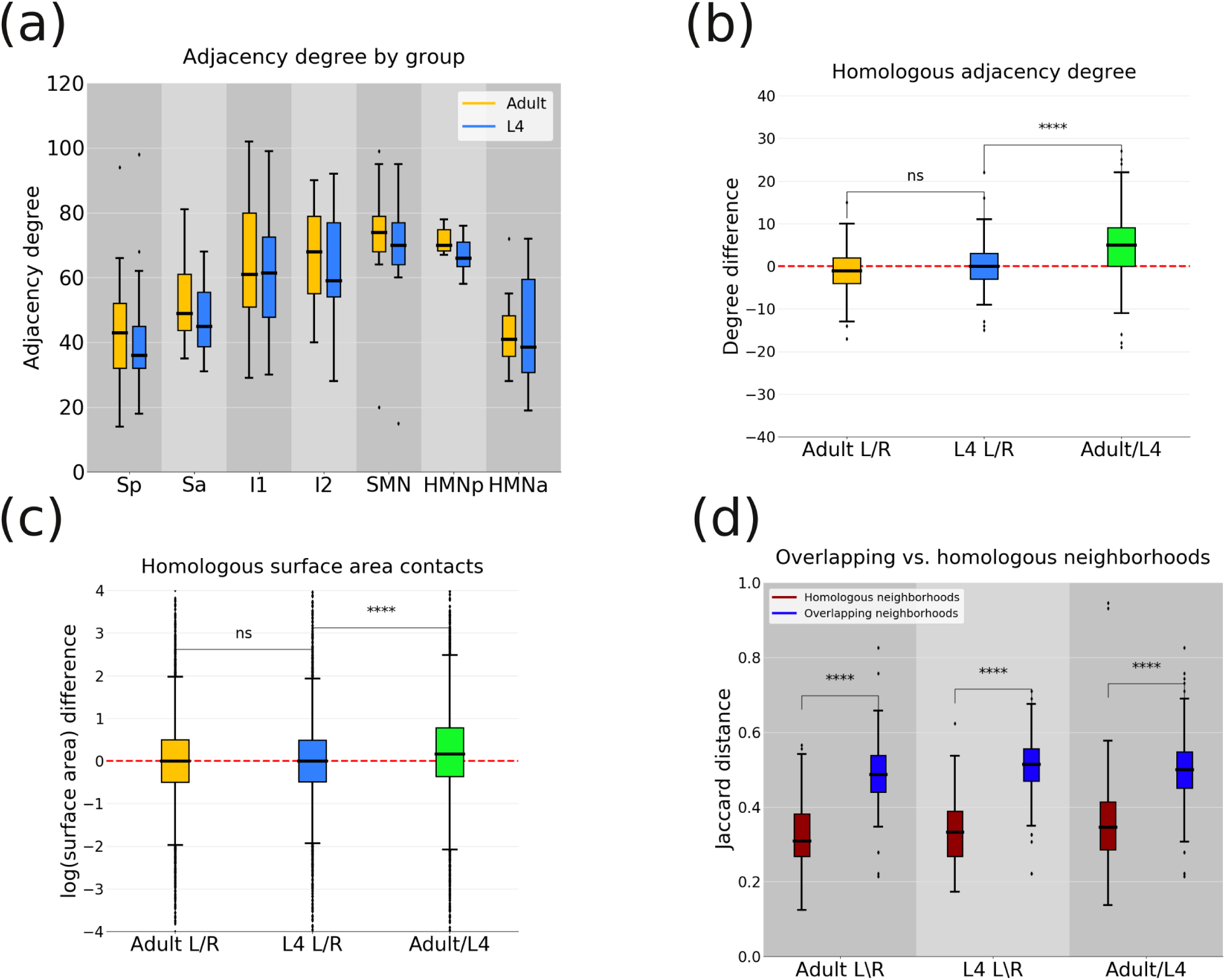
Variability in adjacency degree and contact. (a) Adjacency degree distributions for the seven anatomical groups (see Table S2). (b) Distribution of degree differences for homologous neurons. Compared contralateral neurons in the adult (Adult L/R), the L4 (L4 L/R) and homologous adult and L4 neurons. (c) Distribution of the differences of the log of adjacency contacts between homologous neurons. (d) Distributions of Jaccard distances for overlapping ipsilateral neighborhoods (blue) and homologous contralateral neighborhoods (red) in the adult and L4. (ns) not statistically significant. (****) *p* < 0.01, *t*-test.

To estimate the specificity in adjacency, we exploit the bilateral symmetry of the worm (see Methods) and compute adjacency differences between contralateral homologous (left-right) neurons (see Methods). We first computed differences in neighborhood sizes between homologous neurons in the L4 and in the adult, and then compared the neighborhood sizes of equivalent neurons between the L4 and adult. Both the L4 and adult exhibit strong bilateral symmetry in neighborhood sizes (paired *t*-test, *p* < 0.01, Figure 3(b-c)). In contrast, adult neurons have slightly larger neighborhoods than the equivalent L4 neurons. While slight and while the comparison is across different individual animals, the neighborhood size increase is statistically significant (e.g., as compared to the differences across contralateral homologous pairs in the L4 and adult, *t*-test, *p* < 0.01). This result appears to suggest that neurons increase neighborhood sizes during juvenile development.

The high bilateral symmetry of the worm suggests that contralateral homologous neurons are similarly positioned to ensure similar neighborhoods and hence similar connectivity. However, similar neighborhood sizes do not necessarily imply similar identities of the neighbors. We asked (i) what is the overlap in the set of neighbors of two homologous neurons and (ii) is this overlap distinctive for specific homologous pairs. Both questions are well addressed with the Jaccard distance, which measures the dissimilarity between two sets and ranges from 0 (for equivalent) to 1 (for completely distinct) sets (see Methods). To address the first question we computed the Jaccard distance between neighborhoods of contralateral homologous neurons (*J_h_*). To address the second question, we compared the contralateral distance with the distance between pairs of ipsilateral neurons: for each neuron, we computed the Jaccard distance (*J_o_*) between the neuron’s neighborhood and the most similar ipsilateral overlapping neighborhood (see Methods). We reasoned that similarity between two arbitrary neighborhoods could arise naturally if multiple neuron pairs shared a common neighborhood. The consequence of such extensive neighborhood overlap could be that interchanging the physical locations of the two neurons would not affect their profile of synaptic connectivity. Therefore, the ipsilateral Jaccard distance provides a benchmark for the distinctiveness of different neighborhoods.

We find a high level of equivalence between the neighborhoods of contralateral homologous pairs of neurons, as compared with ipsilateral neurons with overlapping neighborhoods (Figure 3(d)). In each of the three test cases (L4 homologous pairs, adult homologous pairs and L4/adult equivalent neurons) the mean ipsilateral dissimilarity is larger than the mean contralateral homologous dissimilarity (*t*-test, *p* < 0.01). From this we conclude that there exists a high level of neuronal spatial specificity leading, on the one hand, to distinct adjacency profiles between neurons in a common neighborhood, and on the other, to highly similar neighborhoods of homologous contralateral neuron pairs. The distinctive neighborhood associated with each neuron class is a strong indicator that relative process placement is highly specified.

Finally, we asked whether the specificity in neuronal processes and corresponding similarity in neuronal adjacency profiles extend to quantitative measures such as the contact area between pairs of neurons. We define the adjacency contact as the amount of membrane contact between two cells *i* and *j*, measured in *μ*m^2^. We compared the adjacency contact between homologous pairs of adjacent cells (e.g for ASH and AVA, we compare ASHL-AVAL contact with ASHR-AVAR contact). We find remarkably similar average contact areas between contralateral homologs, both in the adult and in the L4 animal (paired *t*-test, *p* < 0.01, Figure 3 (c)). Consistent with the growth in the neighborhood size over development, we also find slightly increased adjacency contacts in adult neurons as compared to the equivalent L4 neurons. Thus, we find that the spatial specificity of neuronal placement gives rise to reproducible contralateral structures, with reproducible relative placement of neuronal processes and even reproducible adjacency contact areas.

### Connectivity fractions vary across neuron classes and synapse type

We next assessed the connectivity fraction of the nerve ring, defined as the ratio of actual to potential synaptic connectivity. The connectivity fraction represents the likelihood that an actual synaptic connection is present at a potential synaptic site (Escobar et al., 2008). This likelihood has become a useful measure of structural plasticity (Escobar et al., 2008; Stepanyants et al., 2002) and may be relevant for information storage within a neural tissue (Chklovskii et al., 2004). For a given neuron, we define the connectivity fraction as the number of synaptic connections divided by the number of neighbors. Because *C. elegans* neurons only make *en passant* synapses, every adjacent neighbor represents a potential synaptic connection. It was previously reported that *C. elegans* neurons make a synaptic connection (presynaptic, postsynaptic or gap junction) with a little more than half of their neighbors (White et al., 1983; White, 1985). Even with the updated wiring diagrams (Cook et al., 2018), we also find a connectivity fraction of ~0.5 when all synaptic connections are considered. However, there is significant variation in connectivity fractions among different synapse types and cell groups which has not been previously reported.

We find that there is no “representative” pre- or postsynaptic connectivity fraction for the nerve ring, because different neurons exhibit different connectivity trends. We distinguish between connectivity fractions for gap junction (*C*^gap^), presynaptic (*C*^Pre^) and postsynaptic (*C*^Post^) connections. *C*^Pre^ and *C*^Post^ vary across sensory, inter- and motor neuron functional classes (Table S2, Figure 4). Not surprisingly, sensory neurons have the highest fractional output (median *C*^Pre^ = 0.28) but the lowest fractional input (median *C*^Post^ = 0.18). The latter reflects considerable lateral sensory-sensory neurons synapses and some feedback connections from first-layer interneurons (Jarrell et al., 2012; Cook et al., 2018). Conversely, motor neurons have the lowest fractional output (median *C*^Pre^ = 0.15) but the highest fractional input (median *C*^Post^ = 0.25). The range of connectivity fractions for interneurons is large (0.5 for both *C*^Pre^ and *C*^Post^) which suggests the mean connectivity fractions are not representative of all interneurons. It is worth noting that a significant fraction of motor neuron output and some sensory/interneuron output is onto head and body wall muscles which are not included in our segmentation. We have estimated what the connectivity fractions would be if muscles were included (Figure S4). While the average connectivity fractions would increase, the variability in connectivity fractions would not substantially change. Therefore, we conclude that pre- and postsynaptic connectivity fractions vary significantly across the nerve ring.

**Figure 4.**
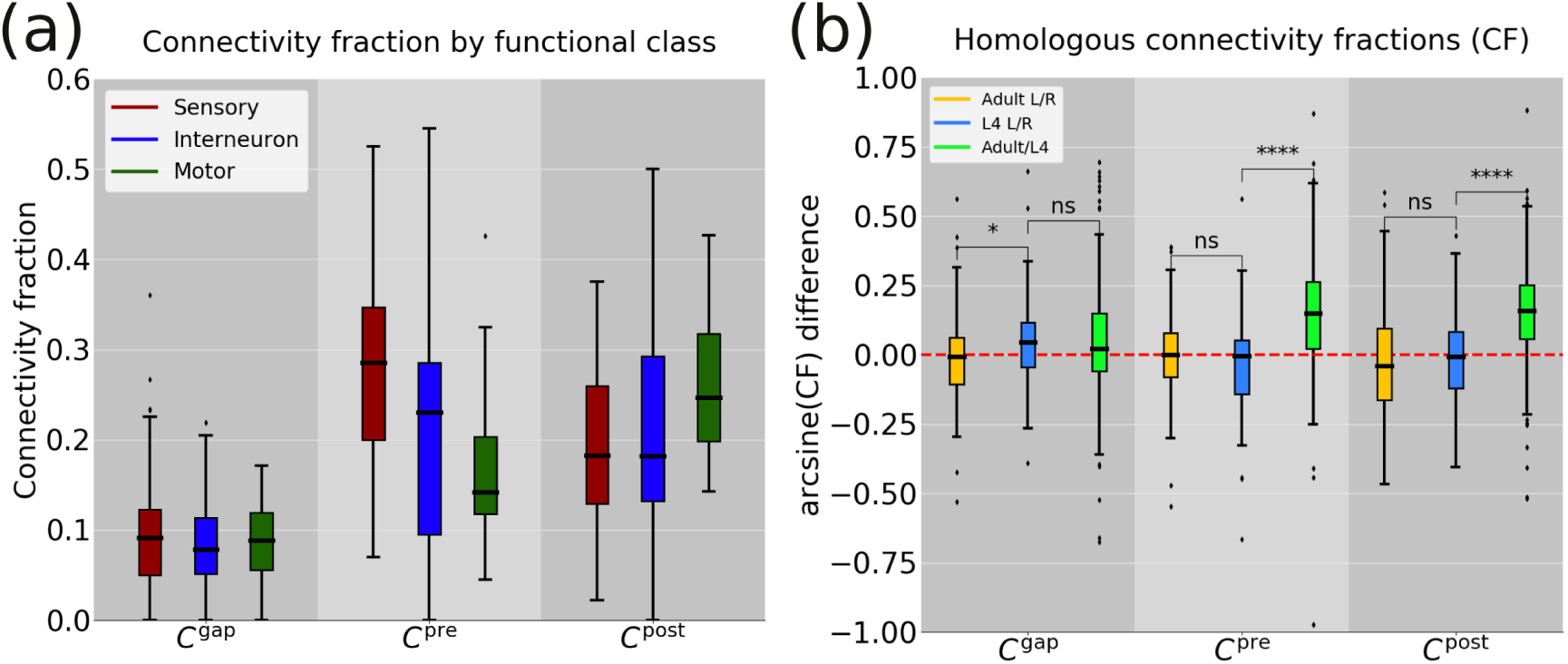
Variability in connectivity fraction. (a) Distribution of connectivity fractions for functional neuron classes. (b) Distribution of the differences between the arcsine of connectivity fractions of homologous neurons. (ns) not statistically significant. (***) *p* < 0.05, *t*-test.

We find that the gap junction connectivity fractions are steady across functional classes and larval development. The mean *C*^gap^ is approximately 0.1 for all functional classes (Figure 4(a)). Unlike *C*^Pre^ and *C*^Post^, we find that (arcsine) *C*^gap^ differences between L4 and adult neurons is not significantly greater than (arcsine) *C*^gap^ differences between contralateral homologous neurons (*t*-test, *p >* 0.05 Figure 4(b)). Collectively, these results suggest pre- and postsynaptic connectivity is class dependent and increases with developmental stage. By comparison, gap junction connectivity is well defined and maintained throughout larval development. This may suggest that chemical and gap junction connectivity serve distinct developmental purposes.

### Synaptic specificity is not a consequence of spatial specificity

We next asked if the reproducibility of synaptic connectivity is due to the reproducibility of process placement. Most synaptic connections in the nerve ring are reproducible between contralateral homologs (Figures 5(a) and S3(a-b)). If synaptic connectivity is completely due to conserved process placement, then the reproducibility of synaptic partners could be described by a purely statistical model, e.g. Peters’ rule (Rees et al., 2017). Under this model, synaptic connections are made with some (perhaps cell autonomous) probability irrespective of neighboring cells. We conclude that such a statistical model is false due to the following four reasons.

**Figure 5.**
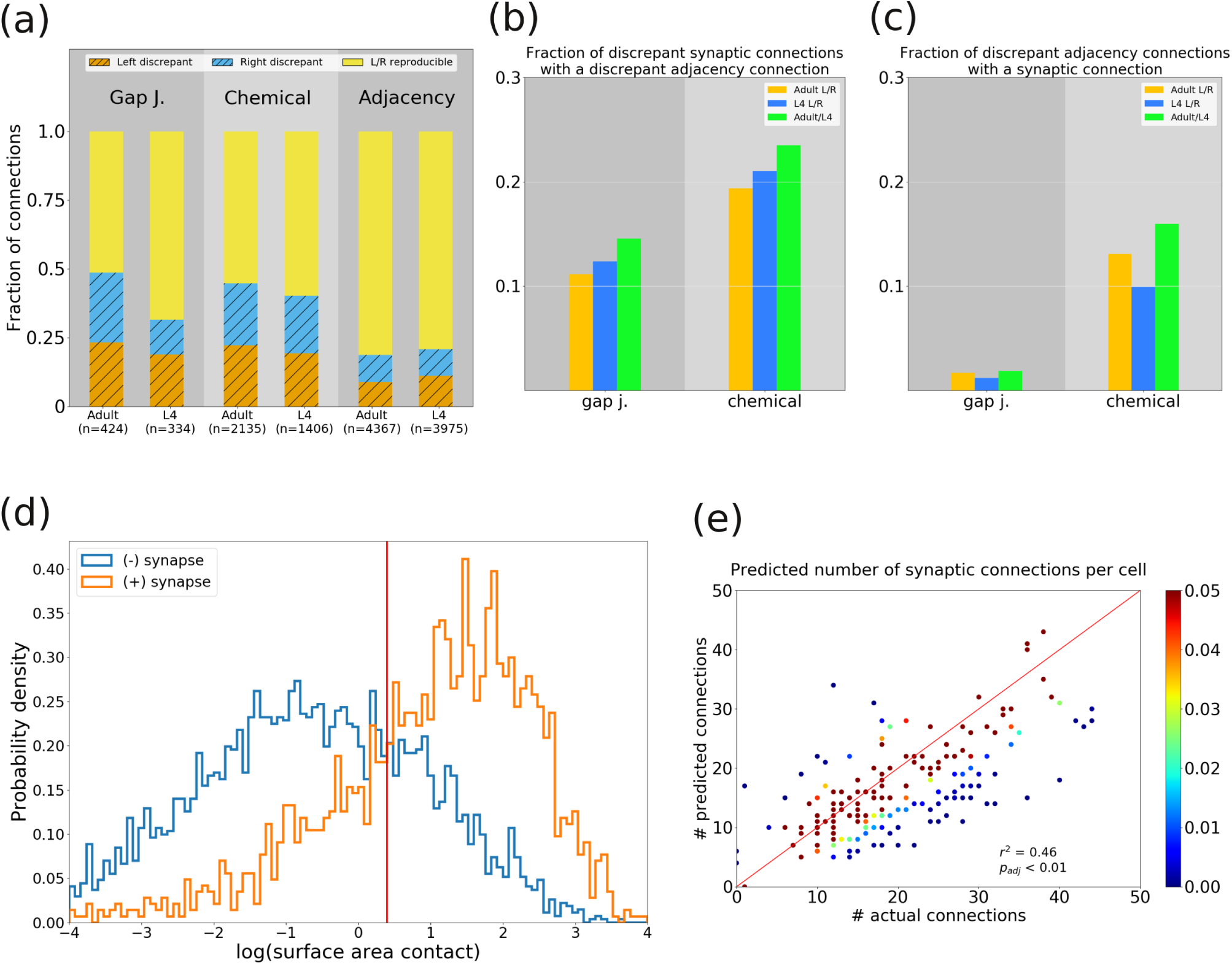
Adjacency variability contributes little to synaptic variability. (a) Fraction of connections that are discrepant. Connections are classified as left discrepant (found only on the left side), right discrepant (found only on the right side) or left/right reproducible (found on both sides of the animal). Numbers represent the number of total connections. (b) Fraction of discrepant synaptic connections that occur at discrepant adjacency connections. (c) Fraction of discrepant adjacency connections with a synaptic connections. (d) Probability density of the log of surface area contacts for adjacencies that do (+) and do not (-) produce a synapse. Red line indicates a decision boundary. Right of the boundary an adjacency has a higher probability of producing a synapse. (e) Predicted number of synaptic connections for each cell compared to the actual number. Predictions made using a logistic regression classifier model. Red line indicates perfect agreement between predicted and actual values. The residual is the distance from the data point to the line. Colors indicate the probability of observing a residual as large or larger. *p*_adj_ is a representative probability for all data points, computed using multiple hypothesis testing.

First, variability in adjacency accounts for less than 20% of the synaptic variability. We say that a connection is discrepant if it occurs on the left (right) side of the worm but is absent on the opposing side. We find that 40–50% of synaptic connections and ~20% of adjacency connections are discrepant (Figure 5(a) and S3(a)). Could the discrepant synaptic connections be due to discrepant process placement of left/right neurons? Roughly 20% of the discrepant synaptic connections occur at discrepant adjacency connections (Figure 5(b)), which suggests that only a small fraction of discrepant synaptic connections could be attributed to differences in process placement. Furthermore, less than 15% of discrepant adjacency contacts yield a synaptic contact (Figure 5(c)), which shows only a small fraction of discrepant adjacency connection contribute to the synaptic connectivity. We also find little correlation between adjacency and synaptic degree differences (*r*^2^ < 0.15, Figure S3(c)). Collectively, these results indicate that only a small fraction of the variability in synaptic connectivity can be attributed to differences in process placement.

Second, synaptic connections are linked to higher adjacency contact, but adjacency contact does not predict connectivity. There is a clear positive relationship between adjacency contact and synaptic probability (Figure 5(d)). Consistent with previous reports (Durbin, 1987), we observe that the (log) distribution of adjacency contacts that do not produce a synapse are skewed to lower surface areas, while the (log) distribution of adjacency contacts that do produce a synapse is skewed towards higher surface areas. To test the predictive power of adjacency contact, we applied a logistic regression classifier (LRC). The LRC is able to predict overall synaptic connectivity with 76% accuracy, but poorly predicts the number of synaptic partners for each cell (Figure 5(e), see Methods). This indicates that adjacency contact is necessary but not sufficient to determine synapse probability.

Third, there are more reproducible synaptic connections than would be expected by chance if synaptic probabilities were held constant among homologous neurons. We constructed a measure called the specificity probability (*p_s_*), defined as the likelihood homologous neurons randomly make the same synaptic contacts among their shared neighbors (see Methods and Figure S5(a)). We say that the likelihood of synaptic reproducibility is low if *p_s_ <* 0.05. We computed specificity probabilities for gap junctions 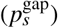, presynaptic 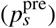 and postsynaptic 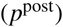 connections. Roughly 30% of cells exhibit a low *p_s_* for all three synapse types, ~45% of cells exhibit a low *p_s_* for two synapse types, ~20% of cells exhibit low *p_s_* for only one of the synapse types and ~5% of cells do not exhibit a low *p_s_* for any synapse type (Figure 6(a)). Therefore, almost all cells (>95%) exhibit some type of connectivity that cannot be accounted for by randomly choosing synaptic connections from a shared set of neighbors. Thus, it seems unlikely that the patterns of synaptic connectivity could be entirely due to random connections among shared neighbors.

**Figure 6.**
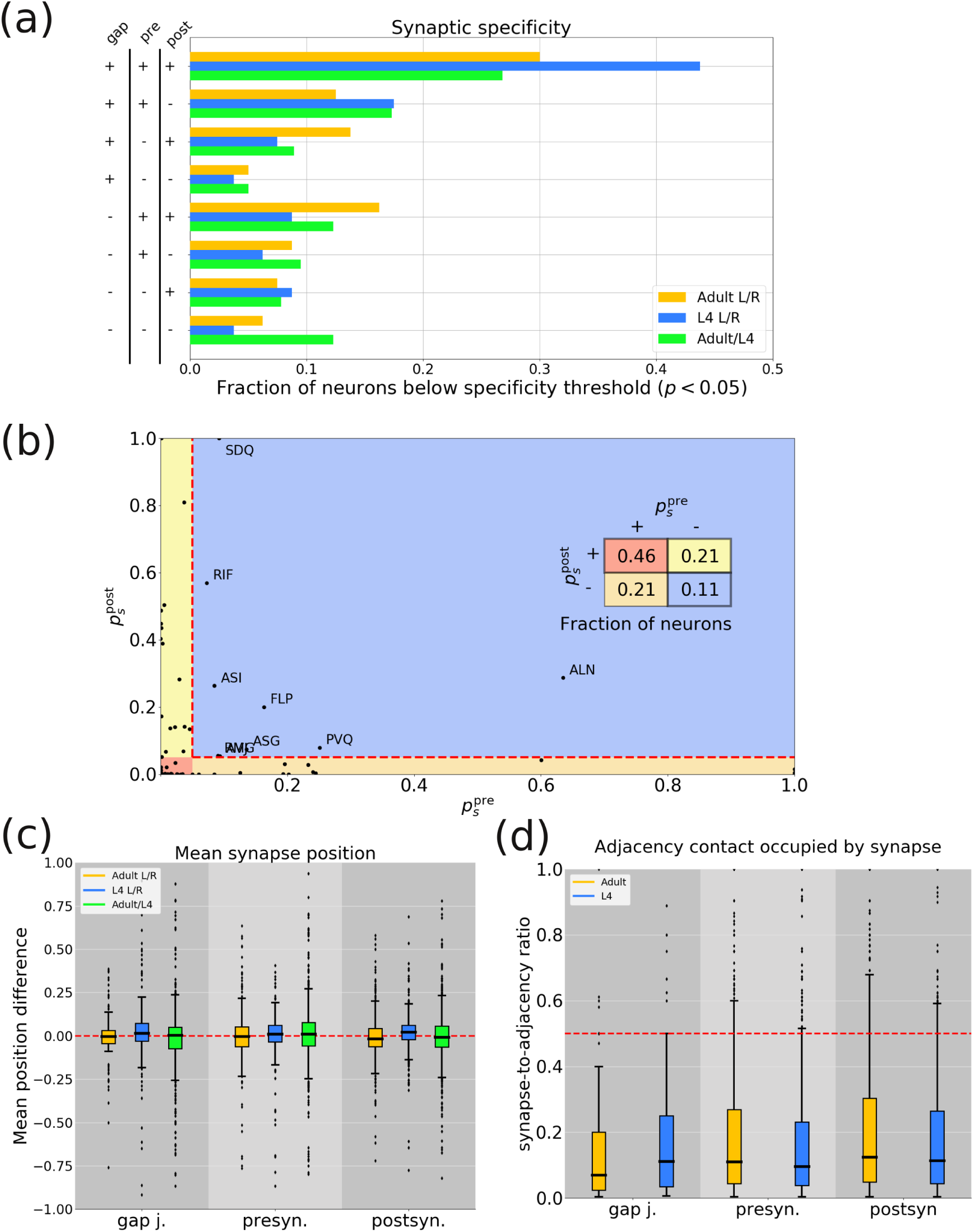
A strict probabilistic model does not account for synaptic reproducibility. (a) Synapse probabilities are either below (+) or above (-) the 0.05 threshold. Left shows the different +/-combinations for 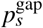, 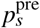 and 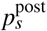. Bar plots show fractions of neurons with the indicated combination of specificity probabilities. Bar color indicates either adult (yellow) or L4 (blue) left/right comparison or comparison between adult and L4 (green). (b) Plot of 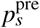 vs. 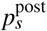 in the adult. Cells fall into one of four categories: (+,+), (+,-), (+,-) or (-,-) indicated by red, yellow, orange and blue, respectively. Table gives the fraction of neurons in each category. Outlier neurons in the last category (-,-) are labeled. Homologous neurons are considered outliers if both 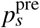 and 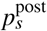 are greater than *α* = 0.05. Red dashed line marks where the probability is 0.05. (c) Distribution of differences between homologous mean synapse positions. (d) Distribution of the synapse-to-adjacency ratios for the adult and L4. Most synaptic contacts are less than 0.5 (red dashed line).

Finally, synapses form at reproducible locations along the neurite despite occupying a small fraction of the surface area contact between cells. We mapped synapses to points along the medial line of each neuron volume and defined the synapse position as the distance from the cell body normalized by the length of the neuron (Figure 6(b), see Methods). In cases where multiple synapses exist between neurons, we took the mean synapse position. We differentiate between gap junction, presynaptic and postsynaptic mean synapse positions. We find that the average difference between homologous mean synapse positions is insignificant (paired *t*-test, *p* > 0.05) but differences can be as high as 25% of the cell length (Figure 6(c)). This suggests that synapse positions are well defined with less than 25% variability. We considered the possibility that adjacency constraints between cells forces synapses to cluster at specific positions, which would suggest that synapse positions are due to the spatial placement of cells. However, we find that the vast majority of synaptic contacts (> 95%) occupy less than half of the surface area between adjacent cells (synapse-to-adjacency ratio, Figure 6(d)). In order for adjacency to account for the 25% variability in synapse position then, we should expect synapse-to-adjacency ratios to be around 0.75. Therefore, we conclude that any constraints on adjacency contact cannot account for the subcellular specificity of synapse positions.

### Synaptic reproducibility is consistent with a combinatorial genetic model

We next assessed if a combinatorial genetic model could account for the reproducibility of synaptic connectivity in the nerve ring. A popular model is the “area code hypothesis”,(Dreyer, 1998) which states that unique cell labels are created by the combinatorial expression of a small-number of cell surface molecules.(Baier, 2013) The observation that some CAMs are differentially expressed and alternatively spliced among neuron populations (Südhof, 2017; Takeichi, 2007; Yagi, 2012; Wojtowicz et al., 2007) seems to support such a hypothesis. To test the hypothesis, we have constructed three different variations of a combinatorial expression model (whole-cell binary expression, subcellular binary expression and isoform expression) and then used curated data of 35 CAM genes with well characterized nerve ring expression (Wormbase WS259 (Harris et al., 2010) and Table S4) to evaluate each model’s ability to uniquely label synaptic partners. We sought to keep our analysis as conservative as possible and therefore only considered CAM genes with well characterized expression (see Methods). That said, the 35 CAM genes can theoretically encode more than 30 billion (2^35^) CAM expression patterns which is more than sufficient to uniquely label the 180 neurons in the nerve ring.

All CAM expression models assume that each neuron is labeled by a combination of CAM proteins (Figure 7(a-b)), but each model assumes a different pattern of CAM expression (see Methods). To evaluate each model, we defined the local uniqueness score (LUS) which is a measure of how frequently a presynaptic neuron can differentiate postsynaptic and nonsynaptic neighbors strictly based on patterns of CAM expression (see Methods). If LUS = 1, then the CAM expression patterns of postsynaptic and nonsynaptic neighbors are always distinct. If LUS = 0, then the CAM expression patterns of postsynaptic and nonsynaptic neighbors are always equivalent. The best CAM expression models should have an average LUS value closer to 1, which would indicate that the model is able to uniquely label synaptic partners for the purposes of wiring specificity.

**Figure 7.**
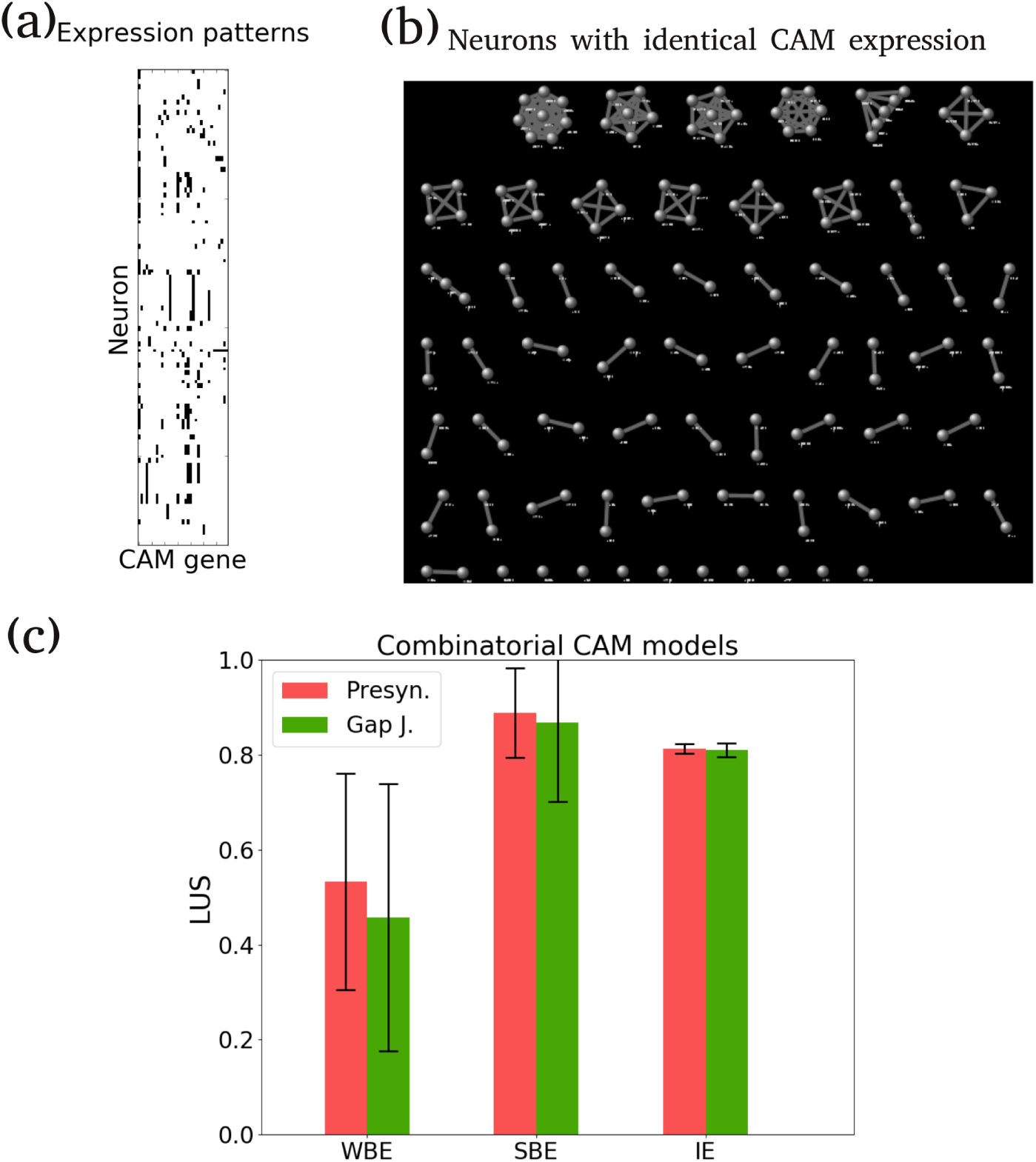
Synaptic reproducibility is consistent with combinatoric genetic models. (a) Expression matrix of 35 CAM genes used in this study. (b) Graph showing neurons (nodes) with identical CAM expression patterns (edges). Clusters indicate that an identical CAM expression is shared by multiple neurons. All clusters contain at least one pair of contralateral homologous neurons. Isolated nodes indicate that the neuron has a unique CAM expression pattern. (c)The LUS scores for the combinatoric CAM models WBE, SBE and IE. Bar heights are the average LUS across neurons. Error bars represent the standard error.

We first consider a minimal, whole-cell binary CAM expression model (WBE) which assumes that CAM protein expression is uniform across the entire cell membrane and that genes are not alternatively spliced (see Methods and Figure S8(a)). We find that this model can only account for about half of the wiring specificity. Under the WBE model, there are only 64 distinct CAM expression labels (the node clusters in Figure 7(b)) and only 10 neuron classes have unique CAM expression labels (the isolated nodes in Figure 7(b)). The mean LUS is 0.55 indicating that only half of postsynaptic partners are distinguishable from nonsynaptic neighbors strictly based on CAM expression (Figure 7(c)). For the given CAM expression data, we conclude that the WBE is not consistent with the reproducibility of synaptic connectivity in the nerve ring.

In contrast, we find that a localized binary CAM expression model can account for ~ 90% of the wiring specificity. The subcellular binary expression (SBE) model assumes that CAM genes proteins are differentially expressed across the cell membrane. The SBE model builds on the previous result that mean synapse positions are conserved and assumes that synapses are created at points of localized CAM expression along the neurite (see Methods and Figure S8(b)). In this model, the presynaptic neuron only needs to distinguish postsynaptic and nonsysnaptic CAM expression labels at points of synaptic contact (Figure S8(b), red ‘x’). The mean LUS score for the SBE model is 0.88 (Figure 7(c)) indicating that comparing expression labels locally at synapses yields a higher frequency of postsynaptic neurons being distinguished from nonsynaptic neighbors. Thus, the additional assumption that CAM proteins are locally expressed along the neurite yields a model that captures more of the synaptic specificity in the network. Further work would be necessary to test to what extent combinatorial CAM expression is localized at points along the neurite and, if so, to what extent such localized CAM expression labels are used to target specific synaptic partners.

In a different whole cell model, we find that an alternatively spliced CAM expression model can account for ~ 85% of the wiring specificity. The isoform expression (IE) model assumes that CAM genes are alternatively spliced and stochastically expressed across nerve ring neurons (see Methods and S9). Alternative splicing allows for a gene to code for multiple isoform proteins and could increase the number of unique CAM expression labels. C. *elegans* exhibits little isoform diversity compared to other organisms (25% of protein-coding genes in C. *elegans* exhibit alternative splicing (Wani & Kuroyanagi, 2017) compared to 95% in humans (Pan et al., 2008)). Moreover, there are rarely more than 10 isoforms expressed by an alternatively spliced CAM gene, compared to the thousands to tens-of-thousands expressed in other organisms (Zipursky & Sanes, 2010). However, the C. *elegans* nervous system may not require such isoform diversity due to its small size. In the nerve ring, there are 15 CAM genes with known alternative splicing which can code up to 9 isoforms (Figure S9). Unfortunately, precise isoform expression of CAM genes in nerve ring neurons is not generally known. Instead, we simulated alternative splicing by randomly assigning a single splice variant for each alternatively spliced gene (see Methods). The average LUS score for the IE model is 0.85 (Figure 7(c)) indicating that isoform expression potentially yields a larger number of unique CAM labels. Based on these simulations, there are on average 140 unique expression patterns when alternative splices are randomly assigned resulting in 107 uniquely labeled neuron classes (Figure S9). Notably, the vast majority of neuron classes express at least one CAM gene with a splice variant (Table S6). This alternative splicing could, in principle, allow neurons to generate unique CAM labels, even without subcellular expression profiles.

## Discussion

To what extent is wiring specificity informed by the spatial proximity of neurons? Clearly spatial factors play a major role in nervous system development, because only adjacent neurons can eventually form a synapse. However, if the relative spatial placement of neurons is well specified, then additional mechanisms may not be required to specify synaptic connectivity. Under these spatial conditions, synapses could form randomly between the neurons and still yield reproducible patterns of connectivity. We explored this question in the *C. elegans* nerve ring, and found that the spatial specificity of the nerve ring cannot fully account for the reproducibility of synaptic connectivity. This suggests that in *C. elegans* both spatial and non-spatial factors contribute to wiring specificity.

We show that the nerve ring aggregates functionally similar synapses and physically segregates distinct computational pathways. The nerve ring neurons can be divided into 7 groups based on function and anatomical location. Synapses between these groups aggregate into radial and azimuthal quasi-layers within the nerve ring. The aggregation of functionally similar synapses to restricted anatomical regions has become a hallmark feature of the spatial specificity of nervous systems across species. For example, the retina has six main cell types whose cell bodies are distributed across three lamina that are in turn coupled by two lamina of synapses.(Baier, 2013) However, unlike the lamina of the retina, the nerve ring quasi-layers are not as well defined. The relatively simple morphology of *C. elegans* neurons likely precludes the possibility of well defined lamina organization because *C. elegans* neurons typically exhibit little if any branching and only make *en passant* synapses. Nevertheless, we find that the mechanosensory and amphid sensory pathway are physically distinct. Mechanosensory pathways are closer to the pharynx where they can directly innervate motor neurons and head muscle arms. Presumably, this placement reduces the number of processing steps between mechanical stimuli and head response. In contrast, the amphids sensory pathways are spatially organized into sensory and interneuron layers, which is consistent with the computational layers observed in the wiring diagram(White et al., 1986; Varshney et al., 2011; Cook et al., 2018) and confirmed by ablations studies(Gray et al., 2005). These results suggest that the structure of the computational network is at least broadly informed by the spatial organization of the processes within the nerve ring.

The present work is the first to detail the layered structure of the *C. elegans* nerve ring, which could support a hierarchical model for nerve ring development. Interneurons and posterior sensory neurons grow around the motor neurons which innervate the layer adjacent to the pharynx. This could point to a hierarchical model for nerve ring development whereby motor neuron innervation occurs first followed by the remaining neurons. Notably, Rapti et al. (2017) analyzed nerve ring assembly during the comma and 1.5 fold stage of embryo development. They imaged a subset of neurons from the amphid, derid and sublateral commissure as well as a few noncommissural neurons. Of these neurons, Rapti *et al*. observed that the sublateral motor neuron axons (SIAV, SIAD, SIBV, SIBD and SMDD) were the first to enter the nerve ring followed by the remaining cells. Furthermore,the remaining cells exhibited aberrant growth when sublateral cells were ablated, suggesting that the nerve ring is hierarchically assembled. Taken together with our volumetric analysis, one possible explanation for the aberrant growth is that the sublateral motor neurons generate the initial tracks of the nerve ring which helps to guide innervation of later axons processes.

Given that process placement is so highly reproducible, one plausible hypothesis is that the reproducibility of synaptic connectivity is largely due to process placement and that any variability of synaptic formation is due to random connectivity among the set of spatially-specified neighbors. Previous studies have suggested that synapse frequency is indeed correlated to the spatial proximity between neurons. This observation has been referred to as Peters’ rule. While there has never been a clear consensus on how this rule should be applied and evaluated (Rees et al., 2017), one simple interpretation of Peters’ rule is that axons make synapses in direct proportion to the number of proximal dendrite targets (Binzegger et al., 2004; Braitenberg & Schüz, 1998). Recent studies have shown that Peters’ rule is not a good predictor of synaptic connectivity (Kasthuri et al., 2015; Mishchenko et al., 2010; Shepherd et al., 2005), but algorithms using variations of Peters’ rule have been able to simulate and reconstruct synaptic connectivity (Markram et al., 2015; Reimann et al., 2015). Consistent with recent ultrastructural analyses(Mishchenko et al., 2010; Kasthuri et al., 2015; Takemura et al., 2015), we find that less than 20% of the variation in synaptic connectivity can be attributed to variation in adjacency and that adjacency contact is too variable to be a reliable indicator of synapse probability. Finally, we show that a statistical model where synapses are randomly made among common neighbors cannot account for the reproducibility of synaptic contacts between homologous neurons. Taken together, these results point to the existence of one or more cellular mechanisms that mediate specific synaptic partnerships; to reliably predict the pattern of synaptic connectivity, such mechanisms must act within the tight developmental regulation of neuronal morphologies and patterns of spatial proximity of cells.

One which have one pre- and postsynaptic cell, a polyadic synapse has one presynaptic cell and two or more postsynaptic cells (Figures S10(a,b)). Each postsynaptic cell is directly apposed to the presynaptic density, which is why all the cells are scored as postsynaptic partners. Roughly 2/3 of synapses are polyadic (Figure S10(c)) and 90% of synaptic connections in the nerve ring involve at least one polyadic synapse (Figure S10(d)). Many of the polyadic synapses are reproducible in reconstructions of the head, body and tail of the worm (White et al., 1986; Hall & Russell, 1991; Varshney et al., 2011). In the nerve ring, roughly a third of polyadic synapses are conserved between homologous neurons (Figure S10(e)), suggesting that the creation of one synapse could (locally) lead to the creation of other synapses (Hall & Russell, 1991). Lack of independence among synaptic connections could serve to amplify variability and make it challenging to discern any statistical relation between adjacency and synaptic connectivity. Interestingly, this effect of polyadic synaptic connectivity may be present in other organisms. In *Drosophilla* medulla neurons, polyadic synapses have been proposed as a possible source of variation in synaptic connectivity (Takemura et al., 2015). In the mouse neocortex, an appreciable fraction of synapses are also polyadic, and Peters’rule fails to capture the redundancy of synaptic connectivity (Kasthuri et al., 2015).

Finally, we show that reproducibility of synaptic connectivity is consistent with a combinatoric genetic model where synaptic partners are identified based on unique combinations of CAM proteins. Our analysis shows that a simple model where uniform binary CAM expression is assumed (WBE) can only account for roughly 55% of the synaptic specificity in the nerve ring. Given known CAM expression, additional gene expression mechanisms are likely required. We proposed two additional mechanisms: localized CAM expression (SBE) and stochastic isoform expression (IE). Given known CAM expression, both additional mechanisms can account for roughly 90% of synaptic specificity in the nerve ring. The localized CAM expression mechanism is additionally supported by subcellular specificity of synapses in the nerve ring. Furthermore, subcellular specificity has also been observed in the *C. elegans* ventral nerve cord, where synaptic tiling of motor neurons DA8 and DA9 depends on the transmembrane Semaphorins and PLX-1/Plexin (Mizumoto & Shen, 2013). Given the low frequency of *C. elegans* genes that exhibit alternative splicing (Wani & Kuroyanagi, 2017), it is perhaps surprising that the IE model can capture 85% of synaptic specificity in the nerve ring. This could be a consequence of the reduced complexity of the *C. elegans* nervous system relative to other organisms. Alternative splicing has been proposed as a mechanism for wiring specificity in other organisms (Zipursky & Sanes, 2010; Südhof, 2017; Yagi, 2012; Neves et al., 2004), which at least suggests that alternative splicing in the *C. elegans* nerve ring is worth further investigation.

Our results are subject to a number of limitations. First, our segmentation is limited to only neuronal processes and does not include other important structures such as glia which also contribute to nerve ring development (Colón-Ramos et al., 2007; Rapti et al., 2017). Second, the sample size is two worms, which makes it very challenging to distinguish between biological and experimental noise. Each data set is a reasonable size of 181 neurons, but these neurons are drawn from 90 distinct neuron classes, each with their own connectivity and adjacency properties (Figure S2). The variability among neuron classes makes it challenging to interpret statistical tests conducted on the collective population of nerve ring neurons. Therefore, we chose to focus on the differences between homologous neurons which is less variable, an approach similar to a recent ultrastructural study of the *Drosophilla* medulla (Takemura et al., 2015). However, inter-worm variability may be greater than intra-worm variability, in which case our analysis would underestimate the variation in adjacency and synaptic connectivity. We tried to address this by comparing the L4 and adult dataset, but because the worms are different ages, it is difficult to distinguish between developmental differences and inter-worm variability. Finally, the CAM expression data is curated, mostly based on transcription fluorescence reporters (so not necessarily based on endogenous expression) and likely incomplete. With this in mind, we have not attempted to identify specific CAM expression patterns which may or may not be used to uniquely label cells. Instead, we have asked if a prescribed model of CAM expression is consistent with the observed wiring specificity in the nerve ring given known CAM expression. However, our models have not considered any potential patterns of temporal CAM expression which may be important for wiring specificity.

Our analysis suggests that both spatial and cell-specific factors contribute to wiring specificity in the *C. elegans* nervous system. Further studies will be required to determine if this can be generalized to all nervous systems or if it is a specific feature of the *C. elegans* nervous system. Having such a small and compact nervous system (only a few hundred neurons) where synapses must be reproduced at single cell resolution may necessitate multiple levels of wiring specificity. By comparison, organisms with significantly larger nervous systems may only need to reproduce synapses between groups of cell types, where a given cell type may consist of hundreds or thousands of neurons. In this instance, it may be energetically prohibitive to specify synaptic wiring with single cell resolution at such a large scale and it may be more efficient to tightly regulate the spatial placement of cells and then allow synaptic connectivity to proceed in a more statistical fashion. If this is the case, then it will be informative to elucidate the evolutionary divergence for these distinct strategies for wiring specificity.

## Methods

### Electron micrographs (EM) preparation and synaptic connectivity

We used legacy electron micrographs (EM) samples originally prepared by (White et al., 1986). These EMs have since been donated from the MRC/LMB archives to the Hall laboratory. The EMs are now digitized and available at www.wormimage.org. This study uses the N2U and JSH data series taken from an adult and L4 hermaphrodite, respectively. The JSH series extends from just anterior of the nerve ring to the excretory pore. The N2U series is substantially longer, extending from just anterior of the nerve ring to the vulva. We only considered the section of the N2U series that physically corresponds to the JSH series. This resulted in roughly 300 sections in the N2U series compared to 400 sections in the JSH series. In N2U, starting at the ventral nerve cord (just posterior to the nerve ring) only every other EM section was imaged. Additionally, it is speculated that the JSH images may have slightly smaller thickness. To correct for this when making comparisons between the L4 and the adult, data from this region in N2U was scaled by a factor of 2. This correction was only necessary for comparing surface area contacts (Figure 3(c)) and synapse to adjacency ratios (Figure 6(d)). For all other analyses, results were checked for consistency between the L4 and the adult.

The volumetric reconstruction was manually done using TrakEM2 software. (Cardona et al., 2012) The software provides tools to allow the user to segment neurons, track the segments and stores the data in XML format. Measurements of the physical contact between neurons was taken directly from the segmented XML data. Synaptic connectivity from these data series was previously reconstructed,(White et al., 1986; Varshney et al., 2011) but we used the most recent reconstruction(Cook et al., 2018) available at www.wormwiring.org. Unlike the previous data(White et al., 1986; Varshney et al., 2011), the wormwiring data contains the spatial locations of synapses which we could project onto our segmented volumes. This dataset also provides estimates of the volumes of synapses which was used to calculate synapse to adjacency contact ratios. Synapse sizes are estimated by the number of serial EM sections in which the synapse was scored.

### Extracting adjacency data

The algorithm for extracting adjacency works as follows. Consider segment *i* from an EM section which is defined by some set of boundary points *B_i_*. For segment *i*, we identify a set of segments *J* that fall within a search radius that is proportional to the radius of *i*. For each segment *j* ∈ *J*, we do a pairwise comparison of boundary points between *B_i_* and *B_j_*. We count the number *n* of pixel pairs that are less than 10 pixels (~ 50 nm) apart. If *n* > 0, then segments *i* and *j* are labeled as adjacent and *n* is added to the adjacency contact between *i* and *j*. This is repeated for each EM section.

To check the accuracy of the algorithm, two TrakEM2 segmented EM sections were manually scored for adjacent neurons by an expert and compared to the adjacency scored by the algorithm (data not shown). In both cases the algorithm outperformed the manual scorer, recognizing adjacent neurons not identified manually. Any failure of the algorithm to identify adjacent neurons (false negatives) was mostly due to poor manual segmentation. For example, the segmentation of the neuron may not have extended to the cell membrane. There were a small number of cases where the algorithm incorrectly labeled two neurons as adjacent (false positives). This was also due to poor segmentation, where the segmentation extended past the cell membrane. In these cases, the surface area contact scored by the algorithm was small and could be screened out in later analyses by requiring adjacent cells have a minimum contact length. Finally, adjacent cells were previously reported for a small subset of neurons based on a sparse analysis of physical adjacency in the L4.(White et al., 1983) The physically adjacent partners identified by our algorithm match those previously reported. Thus, we conclude that when cells are correctly segmented with TrakEM2, the algorithm adequately identifies all physically adjacent neurons.

### Measuring adjacency variation

We estimate variation in adjacency by exploiting the bilateral symmetry of the worm. The worm is bilaterally symmetric and has homologous contralateral neurons on the left and right side of the animal.(White et al., 1986) With few exceptions (e.g. ASE and AWC), contralateral neuron pairs are genetically, functionally and anatomically identical.

We compute the degree difference between contralateral homologous neurons. To assess if the mean contralateral degree difference is significantly different than 0, we used a paired *t*-test. We justify the use of the paired *t*-test, by the assumption that contralateral homologs are representatives of the same neuron class.(White et al., 1986) To test the significance of the developmental degree difference between the L4 and the adult, we compared the distributions of the developmental and contralateral degree differences using the standard *t*-test.

We compute the contact difference between contralateral homologous neurons. Most adjacency contacts are small, ~50% of contacts are less than 0.35 *μ*m^2^ (Figure S2(b)). Excluding the tails, the log of the adjacency contact distribution corresponds to a normal distribution (Figure S2(c) inset). Therefore, we applied a log transform to the adjacency contacts which made the distribution of the differences in adjacency contact more normal. We then used a paired *t*-test to determine if the contralateral contact differences are significant. As above, we used the standard *t*-test the test the significance of the developmental contact difference between the L4 and the adult.

### Comparing overlapping neighborhoods with contralateral neighborhoods

Two neurons that innervate the nerve ring together are adjacent to many of the same neighbors. Hence, the two neurons are said to have overlapping neighborhoods. Let *N*(*i*) be the set of neighbors for neuron *i* and let neuron *j* ∈ *N*(*i*). Then in practice, it is typically the case that *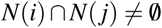*, i.e. the neighborhoods of *i* and *j* overlap. We would like a measure of the difference between two overlapping neighborhoods. A popular metric is the Jaccard distance, which (in this context) is computed as

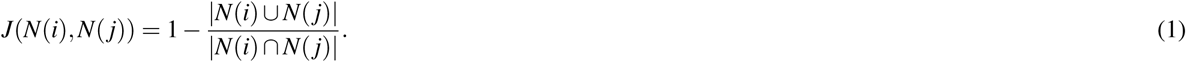

In order to be as conservative as possible, for each neuron *i* we compute the minimum Jaccard distance over its set of neighbors,

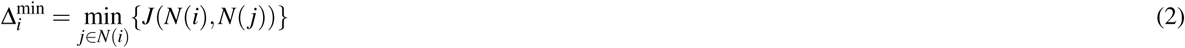

We computed Δ^min^ for each neuron, which gives a distribution of the minimum differences between overlapping neighborhoods. As a control, we computed the Jaccard distance between all homologous neurons within a dataset. Because Jaccard distances are a proportion, we applied an arcsine to both sets of data in order to make the data more normal. We then used a *t*-test to compare both groups.

### Connectivity fraction

The connectivity fraction is the fraction of neighbors that are synaptic partners. Let *d_i_* be the adjacency degree of neuron *i*. Let 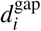, 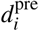 and 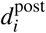 be the number number of gap junction, presynaptic and postsynaptic connections, respectively, of neuron *i*. The pre, post and gap connectivity fractions of neuron *i* are defined as 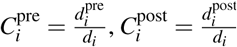 and 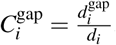, respectively. Because connectivity fractions are a proportion between 0 and 1 and the distributions of connectivity fractions tend to skew to 0, we applied a standard arcsine transformation in order to make the distributions more normal when comparing the connectivity fractions of homologous neurons.

### Logistic Regression Classifier (LRC)

We used the machine learning library scikit-learn(**?**) to build a LRC model for our adjacency data. Once fit to our data, the LRC model classifies adjacency contact between two cells as either a synapse or no synapse. Following Mishchenko *et al*.,(Mishchenko et al., 2010) we assessed the model’s ability to capture variation in synaptic connectivity among neurons by comparing the actual number of synaptic connections for each neuron with the value predicted by the model. Let the random variable *Z_i_ = Y_i_*_1_ *+ Y_i_*_1_ *+*⋯+ *Y_iM_* be the total number of synaptic connections that neuron *i* makes with its *M* neighbors. If synaptic connections are each made independently, then *Y_ij_* has a binomial distribution. Therefore, the expected number of synaptic connections is given by

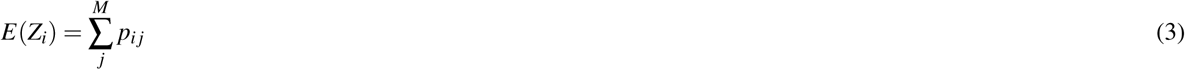

with variance

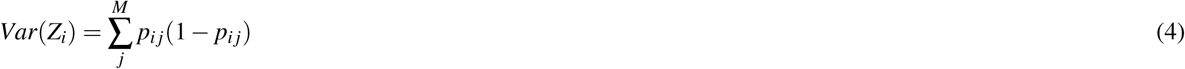

For each neuron, we compute a *p*-value, the probability of observing a discrepancy as great or greater by chance between the actual and expected number of synaptic connections. A representative *p*-value for all the neurons is computed using the Benjamini-Hochberg procedure, (Benjamini & Hochberg, 1995; Benjamini & Yekutieli, 2001) which corrects for the increased chance of observing a Type I error (i.e. falsely rejecting the null hypothesis) due to multiple comparisons and has greater statistical power than the more commonly used Bonferroni correction.(Perneger, 1998) For *m* neurons, the *p* values are arranged in ascending order, *p*_1_ ≤ *p*_2_ ≤ … ≤ *p_m_*, and each *p* value is adjusted to *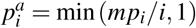*. The multiple hypothesis adjusted *p* value is defined as *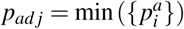*, which is then compared to the false discovery rate *α =* 0.05. When *p_adj_* < 0.05, we reject the null-hypothesis that the model captures the variation in synaptic connectivity.

### Specificity probability

Before proceeding, it is useful to develop some terminology. Contralateral left/right neuron pairs are referred to as homologous neurons. For example, (ASHL,ASHR) and (AVAL,AVAR) are both homologous neuron pairs. Because AVAL and AVAR are physically adjacent neighbors of ASHL and ASHR,respectively, we say that AVA is a homologous neighbor of ASH. Bilaterally conserved synaptic connections are synaptic connections that occur on both the left and right side of the animal. For example, the synaptic connections ASHL→AVAL and ASHR→AVAR are bilaterally conserved connections. We also say that ASH→AVA is a symmetric connection. A synaptic connection that is not bilaterally conserved, i.e. a synaptic connection that occurs on either the left or right side, is said to be an asymmetric connection. An asymmetric connection on the left side is said to be left asymmetric while an asymmetric connection on the right side is said to be right asymmetric.

Let *M* be the number of homologous neighbors, *s* the number of symmetric connections, *a_l_* the number of left asymmetric connections and *a_r_* the number of right asymmetric connections. The left and right connectivity fraction are given by 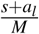 and 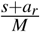, respectively. The left/right connectivity fractions are assumed to be constant while the choice of synaptic partners is random. The number of ways of randomly choosing *s + a_l_* synaptic partners from *M* neighbors is given by the binomial coefficient 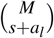. The number of ways of choosing *s* + *a_r_* synaptic partners from *M* neighbors is 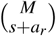. The number of possible combinations between the left and right homologous neuron is given by 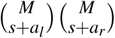. The number of ways of having *s* symmetric connections is given by the multinomial coefficient

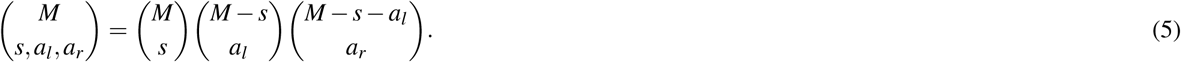

Therefore, the probability of randomly having *s* symmetric connections is given by

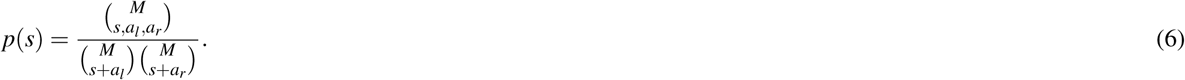

To test the null hypothesis, we need to compute the probability of having *s* or greater symmetric connections. Without loss of generality, assume that *a_l_* ≤ *a_r_*. Then the maximum possible number of symmetric connections is *s* + *a_l_*. Let *k* be a dummy variable such that 0 ≤ *k* ≤ *a_l_*. Note that if the number of symmetric connections is increased to *s* + *k*, then the number of left and right asymmetric connection must be reduced to *a_l_* − *k* and *a_r_* − *k*, respectively. The number of possible ways of having *s* + *k* symmetric connections is given by

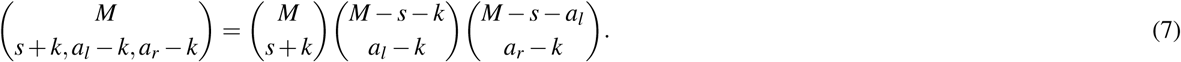

Then the probability of having *s*+ *k* symmetric connections is given by

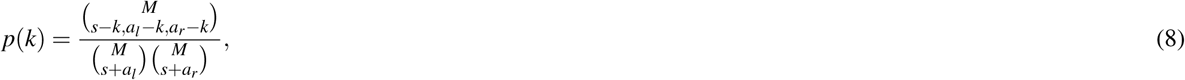

where the probability is a function of *k* and not *s* + *k* because *s* is held constant while *k* is allowed to vary. Finally, the probability of observing at least *s* symmetric connections is given by

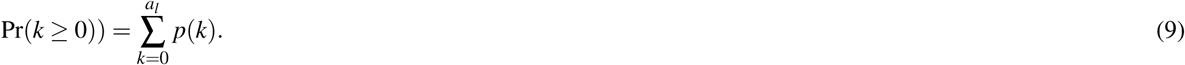

We define the synapse probability as *p_s_ =* Pr(*k* ≥ 0). Using the standard Type I error rate *α =* 0.05, we say that a given pair of homologous neurons exhibit synaptic specificity if Pr(*k* ≥ 0) ≤ 0.05. Here, we have computed the probability of bilaterally conserved presynaptic connections, but the probability of bilaterally conserved postsynaptic connections is computed in a similar way.

If the appropriate substitutions are made, equation (9) can also be used to compute the probability of observing at least *s* developmentally conserved connections. Specifically for a given neuron, let *M* be the number of shared neighbors in both the L4 and the adult, let *a_l_* be the number of synaptic connection in the L4 but not the adult and let *a_r_* be the number of synaptic connections in the adult but not the L4.

### Measuring reproducibility of synapse positions

We mapped synapses to positions along the medial line of the neuron volume. Neuron processes in the nerve ring can mostly be characterized as ribbon like structures that mostly exhibit no branching. Hence, mapping synapse positions to a one dimensional distance along a line is a valid approximation of synapse positions. We applied a mesh contraction algorithm(Au et al., 2008) to the reconstructed volume of each cell in order to get the medial line (skeleton) of the neural process. For each skeleton, we defined the reference point as the point on the skeleton closest to the cell body. We then mapped synapse positions to points on the skeleton and measured the length of the skeleton from the reference point to the synapse position (Figure S6(a)). To facilitate comparisons between cells, we normalized the measured length by the total length of the skeleton. Hence, synapse positions represent the normalized distance that must be traveled from the cell body to the synapse along the neural process.

Because roughly 53% of adult synaptic partners form 2 or more synapses and the number of synapses between neurons varies between homologous neurons, we compared the mean synapse position for homologous synaptic partners It should also be noted that we only compared mean synapse positions of cells that exhibited synaptic specificity (i.e. *p_s_ <* 0.05). If a cell does not exhibit presynaptic specificity and synaptic connections are seemingly random, then it makes little sense to expect mean synapse positions to be conserved (similar reasoning for gap junctions and postsynaptic synapses).

As an example, Table S3 shows the synapses where AIZL and AIZR are presynaptic. The numbers next to the postsynaptic partners are the normalized synapse positions and length of the synapse. The synapse position is the number of EM sections traversed from the reference point to the synapse point. The reference point is the point on the reconstructed skeleton closest to the cell body. The synapse weight is the length of the synapse given as the number of EM sections. The mean synapse position is the average of the individual synapse positions weighted by the synapse weights. The correlation is computed between the weighted mean of the synapse positions. Figure S6 illustrates the correlation between the mean synapse positions for AIZL and AIZR in the adult. For this particular contralateral pair, mean synapse positions for all three synapse types show a strong positive correlation. This process is repeated for all neurons that exhibit synaptic specificity.

### Cell adhesion molecule (CAM) expression in the nerve ring

The *C. elegans* neuronal genome has 106 cell adhesion molecules (CAM) genes(Hobert, 2005) and there are 895 genes expressed in the head neurons (Wormbase WS259 (Harris et al., 2010)). Comparing the two lists, there are 55 CAM genes that are expressed in nerve ring neurons. We designated the nerve ring CAM genes to one of two categories based on how precisely the expression patterns have been characterized: CAM I and CAM II. CAM I consists of 35 genes whose expression in the nerve ring has been clearly identified and linked to specific neurons (Table S4). CAM II consists of 17 genes that are said to be expressed in all nerve ring neurons or for whom subsets of nerve ring neurons are not clearly identified (Table S5). For example, expression of the CAM II gene *egl-15* is observed in hypodermal cells, sex myoblasts, the type I vulva muscles and some “unidentified” head neurons (Huang & Stern, 2004). Because the head neurons were not clearly identified, *egl-15* was placed in CAM II. In order to keep the results as conservative as possible, CAM II neurons were removed from the analysis and only CAM I genes were considered. Henceforth, when CAM genes are mentioned, it should be understood that the CAM I genes are being referenced.

The expression of CAM genes in the nerve ring is sparse with neurons typically expressing a relatively small number of genes and single genes being expressed across multiple neurons. There are 28 nerve ring neurons that have no known CAM expression and were subsequently removed from the analysis. However, most of the remaining neurons express up to 5 CAM genes and over 60% of CAM genes are expressed in at least 5 neurons (Figure S7). Neuron PVT expresses the most CAM genes (11) and gene *cam-1* is expressed in the most neurons (70). There is no discernible structure to the expression matrix (Figure 7a). A number of algorithms were applied to the matrix (e.g. diagonalization and bipartite graph clustering), but none yielded any meaningful organizational insights.

### Calculating local uniqueness score (LUS)

We identified 35 CAM genes with well characterized expression in the nerve ring from which we constructed an expression matrix *E*. Expression patterns were determined from the expression matrix, where *E_ij_* = 1 if neuron *i* expresses gene *j* and *E_ij_* = 0 otherwise (Figure 7a). Hence, the *i*th row of the expression matrix can be used to generate the binary expression label *e_i_* for the *i*th neuron. If *e_i_* = *e_j_* for neurons *i* and *j*, we say that the two neurons have equivalent CAM expression patterns. To evaluate the proposed combinatorial expression models we define the local uniqueness score (LUS).

For the whole-cell binary CAM expression model (WBE) and the isoform expression model (IE), the LUS was computed at the level of synaptic partners (Figure S8(a)). For a given neuron, we determined the CAM expression of the postsynaptic partners and the nonsynaptic physically adjacent neighbors. Let *s* be the number of postsynaptic partners and *m* be the number of postsynaptic partners whose expression matches at least one nonsynaptic neighbor. The LUS is defined as

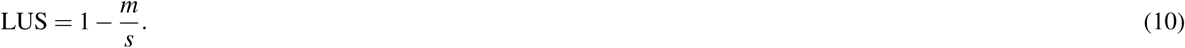

For the subcellular binary expression model (SBE), the LUS was computed at the synapse level (Figure S8(b)). At each synapse for a given neuron, we compared the CAM expression of the postsynaptic neurons and nonsynaptic physically adjacent neighbors. For synapse *i*, let *s_i_* be the number of postsynaptic neurons and *m_i_* be the number of postsynaptic neurons whose expression matches at least one of the physically adjacent neighbors. If the neuron has *M* synapses, the LUS is given by

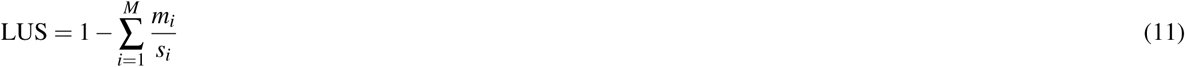

## Acknowledgments

We thank Jonathan Hodgkin and John White for their help in donating archival TEM material from LMB/MRC to the Hall lab for curation. We thank O. Au for generously providing the mesh contraction software. CB was supported by the Leeds International Research Scholarship. DHH is supported by NIH OD 010943. NC was supported by EPSRC EP/J004057/1.

## Author contributions

CB, SC and SE conceived the project. CB and SC segmented the electron micrographs. CB analyzed data. CB and NC drafted the manuscript. DHH curated the data. NC, DHH and SE provided critical revisions. All authors approved final version of the manuscript.

## Supplemental Materials

Prior to publication all supplementary data including raw data can be downloaded at http://wormwiring.org/brittin-etal-2018-data/. All code used for analysis can be accessed at https://github.com/cabrittin/volumetric_analysis.

**Table S1.** Summary of volumeric reconstructions. NB: Volume estimated from the sum of the cross-sectional areas of the segmented neurons.

|  | Adult | L4 |
| --- | --- | --- |
| Series name | N2U | JSH |
| # EMs | 302 | 410 |
| Volume ($\mu\text{m}^3$) | 4051 | 3533 |
| # Neurons | 185 | 181 |
| # adjacency connection | 5368 | 4861 |
| # chemical connections | 2349 | 1583 |
| # chemical synapses | 4173 | 2666 |
| # gap junction connections | 490 | 404 |
| # gap junction synapses | 730 | 702 |

**Table S2.** Functional and anatomical cell groupings.

| Functional | Anatomical | Neurons |
| --- | --- | --- |
| Sensory | Sa | BAGL,BAGR,CEPDL,CEPDR,CEPVL,CEPVR,FLPL,FLPR,IL1DL,IL1DR,IL1L,IL1R,IL1VL,IL1VR,IL2DL,IL2DR,IL2L,IL2R,IL2VL,IL2VR,OLLL,OLLR,OLQDL,OLQDR,OLQVL,OLQVR,URXL,URXR,URYDL,URYDR,URYVL,URYVR |
|  | Sp1 | ASGL,ASGR,AVM,SDQL,SDQR |
|  | Sp2 | ADEL,ADER,ADFL,ADFR,ADLL,ADLR,AFDL,AFDR,ALML,ALMR,ALNL,ALNR,AQR,ASEL,ASER,ASHL,ASHR,ASIL,ASIR,ASJL,ASJR,ASKL,ASKR,AWAL,AWAR,AWBL,AWBR,AWCL,AWCR,DVA,PLNL,PLNR |
| Interneuron | I1 | ADAL,ADAR,AIAL,AIAR,AIBL,AIBR,AIML,AIMR,AINL,AINR,AIYL,AIYR,AIZL,AIZR,ALA,AUAL,AUAR,AVAL,AVAR,AVBL,AVBR,AVDL,AVDR,AVFL,AVFR,AVHL,AVHR,AVJL,AVJR,BDUL,BDUR,PVCL,PVCR,PVNL,PVNR,PVPL,PVPR,PVQL,PVQR,PVR,RICL,RICR,RID,RIFL,RIFR,RIGL,RIGR,RIR,RMGL,RMGR |
|  | I2 | AVEL,AVER,AVKL,AVKR,AVL,DVC,PVT,RIAL,RIAR,RIBL,RIBR,RIH,RIML,RIMR,RIPL,RIPR,RIS,RMFL,RMFR,SAADL,SAADR,SAAVL,SAAVR,URBL,URBR |
| Motor | HMNa | RMED,RMEL,RMER,RMEV,URADL,URADR,URAVL,URAVR |
|  | HMNp | RIVL,RIVR,RMDDL,RMDDR,RMDL,RMDR,RMDVL,RMDVR,RMHL,RMHR |
|  | SMN | SABD,SIADL,SIADR,SI AVL,SI AVR,SIBDL,SIBDR,SIBVL,SIBVR,SMBDL,SMBDR,SMBVL,SMBVR,SMDDL,SMDDR,SMDVL,SMDVR |

**Table S3.** Adult AIZ Synapse postitions. Outside columns give the postsynaptic partners of AIZL and AIZR. The interior columns give the normalized synpase position and synapse weight. Synapse weight is the number of EM sections where the synapse was scored. The center columns give the means of synapse positions weighted by the synapse weights.

| AIZ synapse positions |  |  |  |  |  |
| --- | --- | --- | --- | --- | --- |
| AIZL |  |  | AIZR |  |  |
| Postsynaptic | Synapse position: (position,weight) | weighted mean | weighted mean | Synapse position: (position,weight) | Postsynaptic |
| ADFL | (0.79,2),(0.78,1) | 0.79 | 0.90 | (0.90,1) | ADFR |
| AIAL | (0.62,1),(0.63,1),(0.64,2),(0.65,1),(0.81,1),<br>(0.79,4),(0.78,1),(0.66,1) | 0.72 | 0.79 | (0.79,1),(0.77,1),(0.78,1),(0.80,1),(0.79,1) | AIAR |
| AIBL | (0.64,2),(0.81,1),(0.62,1),(0.79,4),(0.63,1) | 0.72 | 0.78 | (0.77,1),(0.78,1),(0.80,1) | AIBR |
| AIBR | (0.60,5),(0.61,4),(0.64,1),(0.58,2),(0.56,6),<br>(0.54,1),(0.55,1),(0.55,1),(0.56,1),(0.57,1),<br>(0.53,4),(0.96,3),(0.57,2),(0.97,1),(0.59,1),<br>(0.60,2),(0.61,1),(0.96,2),(0.97,1),(0.98,2),<br>(0.99,3),(0.93,1) | 0.69 | 0.62 | (0.90,1),(0.98,2),(0.64,2),(0.64,1),(0.67,2),<br>(0.55,1),(0.56,3),(0.62,2),(0.51,9),(0.59,4),<br>(0.61,4),(0.61,2),(0.96,2),(0.54,3),(0.58,11) | AIBL |
| ASEL | (0.64,2),(0.65,1),(0.66,1) | 0.65 | 0.79 | (0.79,1),(0.79,1) | ASER |
| AVEL | (0.96,3) | 0.96 | 0.94 | (0.94,1) | AVER |
| AVER | (0.59,1),(0.61,1),(0.56,6),(0.61,4),(0.54,1),<br>(0.55,1),(0.56,1),(0.57,1),(0.53,4),(0.64,1),<br>(0.57,2),(0.58,1),(0.58,2) | 0.57 | 0.57 | (0.62,2),(0.64,2),(0.67,2),(0.51,9),(0.61,2),<br>(0.58,11) | AVEL |
| DVA | (0.91,1) | 0.91 | 0.95 | (0.94,1),(0.96,1),(0.94,1) | DVA |
| RIAL | (0.31,2),(0.42,2),(0.53,1),(0.53,2),(0.44,1),<br>(0.55,5),(0.33,4),(0.41,1),(0.29,2),(0.48,3) | 0.43 | 0.44 | (0.42,2),(0.41,1),(0.31,2),(0.44,1),(0.35,4),<br>(0.30,1),(0.51,2),(0.53,1),(0.53,5),(0.54,4),<br>(0.33,2) | RIAR |
| RIML | (0.60,2),(0.61,1),(0.56,6),(0.95,2),(0.96,2),<br>(0.99,3),(0.98,2),(0.60,5),(0.55,1),(0.53,4),<br>(0.57,6),(0.97,1) | 0.68 | 0.65 | (0.62,3),(0.54,3),(0.64,1),(0.55,1),(0.56,3),<br>(0.51,9),(0.59,4),(0.61,4),(0.98,2),(0.96,2),<br>(0.99,1),(0.99,1),(1.00,1) | RIMR |
| SMBDL | (0.34,1),(0.31,2),(0.29,2),(0.42,2),(0.33,2),<br>(0.33,1),(0.41,1),(0.44,1),(0.55,5),(0.45,2),<br>(0.37,1),(0.47,4) | 0.42 | 0.35 | (0.38,3),(0.31,2),(0.33,2),(0.26,4),(0.59,1),<br>(0.53,1),(0.35,4) | SMBDR |
| SMBVL | (0.29,2),(0.33,2),(0.33,1),(0.41,1),(0.45,2),<br>(0.37,1),(0.31,3),(0.47,4),(0.48,3) | 0.40 | 0.37 | (0.38,3),(0.51,2),(0.44,1),(0.26,4) | SMBVR |

**Table S4.** CAM genes with well characterized expression in NR neurons. Pre and post columns indicate whether genes are expressed in the pre- and/or postsynaptic neuron.

| CAM I genes |  |  |  |  |
| --- | --- | --- | --- | --- |
| Protein family | Gene name | Pre | Post | Isoforms |
| Ig domain | *cam-1* | + | + | 3 |
|  | *ver-3* | + | + | 1 |
|  | *igcm-1* | + | + | 1 |
|  | *igcm-2* | + | + | 1 |
|  | *oig-1* | + | + | 1 |
|  | *oig-3* | + | + | 1 |
|  | *rig-1* | + | + | 2 |
|  | *rig-3* | + | + | 1 |
|  | *rig-4* | + | + | 1 |
|  | *rig-5* | + | + | 7 |
|  | *rig-6* | + | + | 4 |
|  | *ncam-1* | + | + | 3 |
|  | *sax-3* | + | + | 2 |
|  | *sax-7* | + | + | 6 |
|  | *lad-2* | + | + | 1 |
|  | *syg-1* | + | + | 2 |
|  | *syg-2* | + | + | 7 |
|  | *unc-40* | + | + | 1 |
|  | *unc-5* | + | + | 6 |
|  | *zig-1* | + | + | 1 |
|  | *zig-2* | + | + | 1 |
|  | *zig-3* | + | + | 1 |
|  | *zig-4* | + | + | 1 |
|  | *zig-5* | + | + | 1 |
|  | *zig-8* | + | + | 1 |
|  | *madd-4* | + | + | 3 |
| Ig + LRR | *pxn-2* | + | + | 1 |
| eLRR | *slt-1* | + | + | 1 |
|  | *tol-1* | + | + | 1 |
|  | *dma-1* | + | + | 1 |
| cadherins | *cdh-3* | + | + | 1 |
|  | *fmi-1* | + | + | 3 |
|  | *casy-1* | + | + | 3 |
| neurexin superfamily | *nlr-1* | + | - | 1 |
| neurexin ligands | *lat-1* | - | + | 3 |
|  | *nlg-1* | - | + | 9 |

**Table S5.** CAM genes that do not have well characterized expression in NR neurons.

| CAM II genes |  |  |  |  |
| --- | --- | --- | --- | --- |
| Protein family | Gene name | Pre | Post | Isoforms |
| Ig domain | *egl-15* | + | + | 16 |
|  | *mig-6* | + | + | 3 |
|  | *oig-2* | + | + | 1 |
|  | *oig-4* | + | + | 1 |
|  | *oig-5* | + | + | 1 |
|  | *zig-6* | + | + | 1 |
|  | *zig-7* | + | + | 1 |
|  | *zig-10* | + | + | 1 |
|  | *igeg-1* | + | + | 2 |
| Ig + LRR | *pxn-1* | + | + | 1 |
|  | *iglr-1* | + | + | 1 |
|  | *iglr-3* | + | + | 2 |
| eLRR | *fshr-1* | + | + | 2 |
|  | *lron-5* | + | + | 1 |
|  | *lron-9* | + | + | 4 |
|  | *lron-14* | + | + | 2 |
| cadherins | *cdh-4* | + | + | 1 |
| neurexin superfamily | *nrx-1* | + | - | 13 |
|  | *bam-2* | + | - | 1 |

**Table S6.** Alternatively spliced CAM genes expressed in bilaterally symmetric neurons.

| Bilaterally expressed alt. spliced genes |  |
| --- | --- |
| Neuron class | Genes |
| ADA | *cam-1,sax-7* |
| ADE | *cam-1* |
| ADF | *syg-1* |
| ADL | *cam-1,syg-1* |
| AIB | *ncam-1,rig-1,rig-6* |
| AIM | *cam-1* |
| AIN | *cam-1,ncam-1,rig-1,rig-5,syg-1* |
| AIY | *cam-1,lat-1,nlg-1* |
| AIZ | *cam-1,madd-4,syg-2* |
| ALM | *cam-1,rig-6,sax-7* |
| ALN | *cam-1,syg-2* |
| ASE | *unc-5* |
| ASG | *madd-4* |
| ASH | *cam-1* |
| ASI | *cam-1,ncam-1* |
| ASJ | *ncam-1* |
| ASK | *cam-1* |
| AUA | *cam-1,rig-1,rig-5,rig-6* |
| AVA | *cam-1,rig-1,rig-6,sax-3* |
| AVB | *cam-1,ncam-1,rig-1,rig-6,sax-3* |
| AVD | *cam-1,rig-1,rig-5,sax-3* |
| AVE | *cam-1,ncam-1,rig-1,rig-6* |
| AVH | *cam-1,madd-4,rig-1,syg-1* |
| AVJ | *cam-1,rig-1* |
| AVK | *cam-1,madd-4* |
| BDU | *cam-1* |
| FLP | *cam-1* |
| HSN | *cam-1,fmi-1,nlg-1,rig-6,sax-3,syg-1* |
| IL1 | *sax-7* |
| IL2 | *sax-7,unc-5* |
| OLL | *casy-1,madd-4,sax-7,unc-5* |
| OLQ | *sax-3,sax-7,unc-5* |
| PLN | *syg-2* |
| PVC | *cam-1,ncam-1,rig-1,rig-6,sax-3* |
| PVP | *fmi-1* |
| PVQ | *cam-1,fmi-1,sax-3* |
| RIA | *madd-4,unc-5* |
| RIB | *rig-6* |
| RIC | *cam-1,madd-4,rig-1,rig-5,rig-6,syg-2* |
| RIF | *rig-5,rig-6,syg-1* |
| RIG | *syg-1* |
| RIM | *cam-1,rig-6,syg-1,syg-2* |
| RIV | *cam-1* |
| RMD | *cam-1,casy-1,rig-1,rig-5,rig-6,sax-3* |
| RME | *cam-1,madd-4,rig-6* |
| RMG | *cam-1,sax-3,sax-7* |
| SAA | *rig-5,rig-6,syg-1* |
| SDQ | *cam-1,fmi-1* |
| SIA | *rig-6,sax-3,syg-1* |
| SIB | *rig-6,sax-3,syg-1* |
| SMD | *casy-1,rig-1,rig-5,rig-6,sax-3* |
| URA | *nlg-1* |
| URB | *nlg-1* |
| URX | *cam-1,rig-6* |

**Figure S1.**
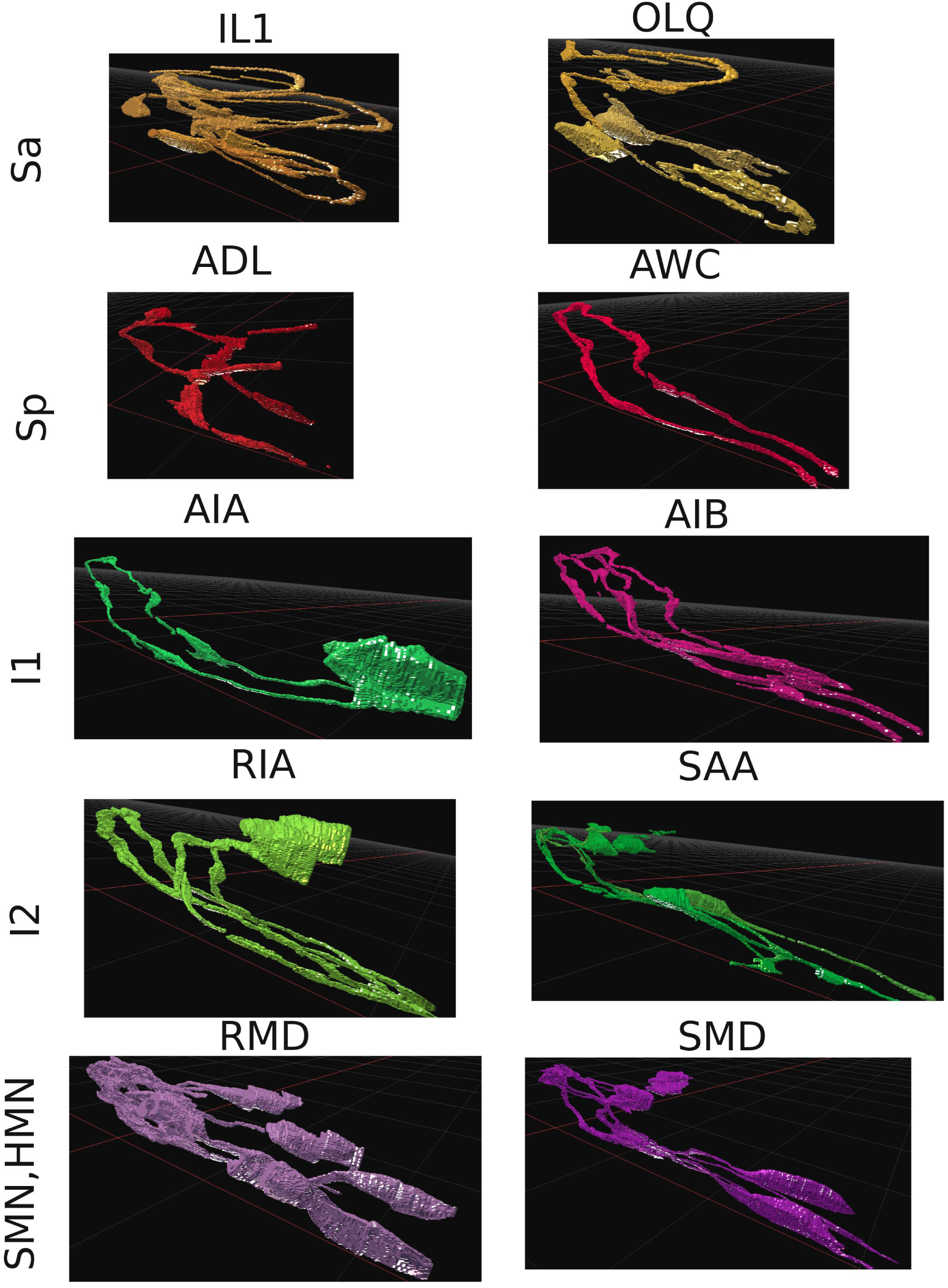
Samples of neuron morphologies. Images take from our web app at http://wormwiring.org/apps/neuronVolume.

**Figure S2.**
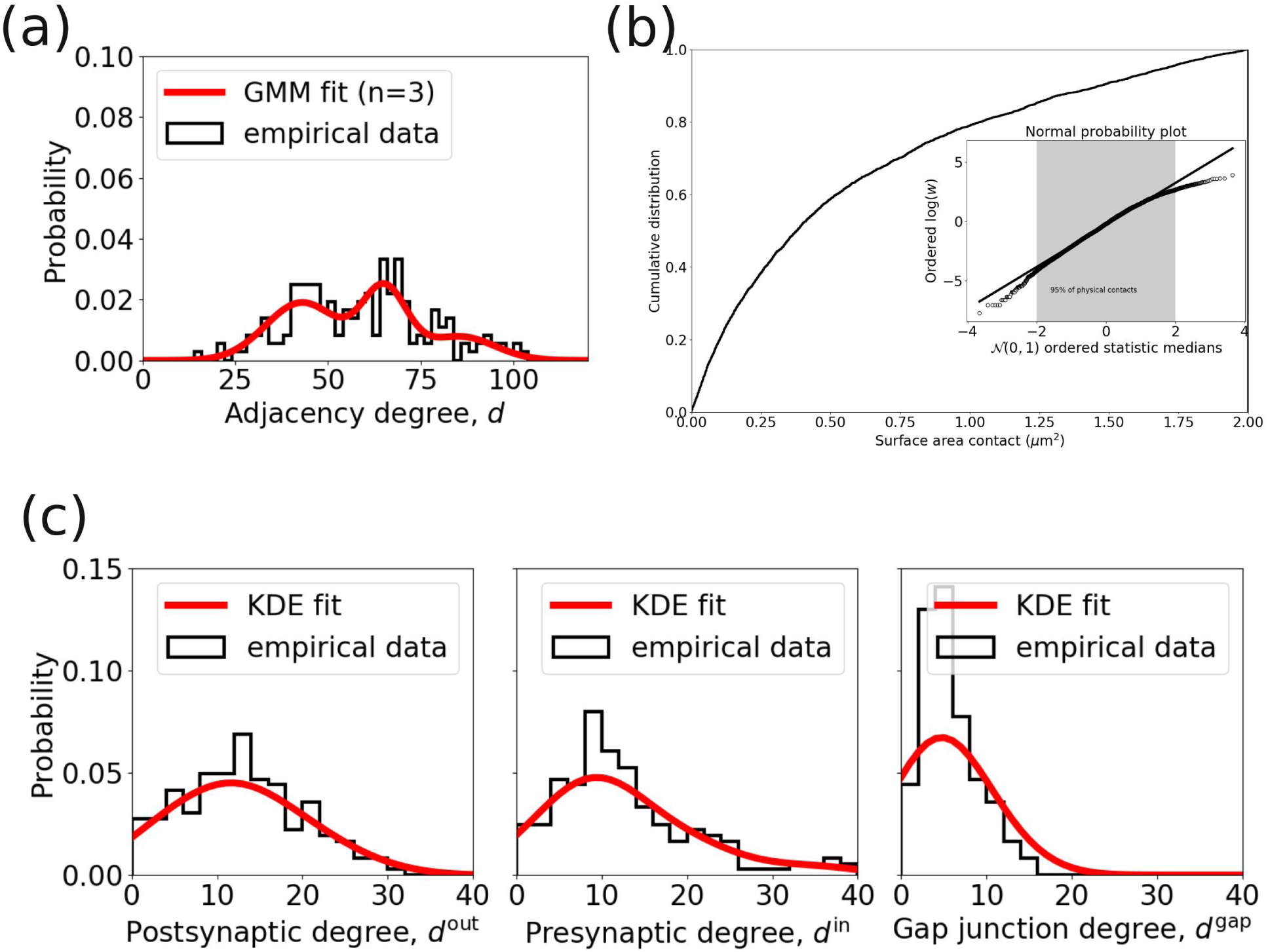
Relevant data distributions. (a) Adult adjacency degree distribution. Fit with a 3 component Gaussian Mixture Model. (b) Cumulative distribution of adult surface area contacts between cells. Inset: Normal probability plot of the log of surface area contacts. The middle ~ 95% of data is lognormal. (c) Adult postsynaptic, presynaptic and gap junction degree distributions. Fit with a kernel density estimator.

**Figure S3.**
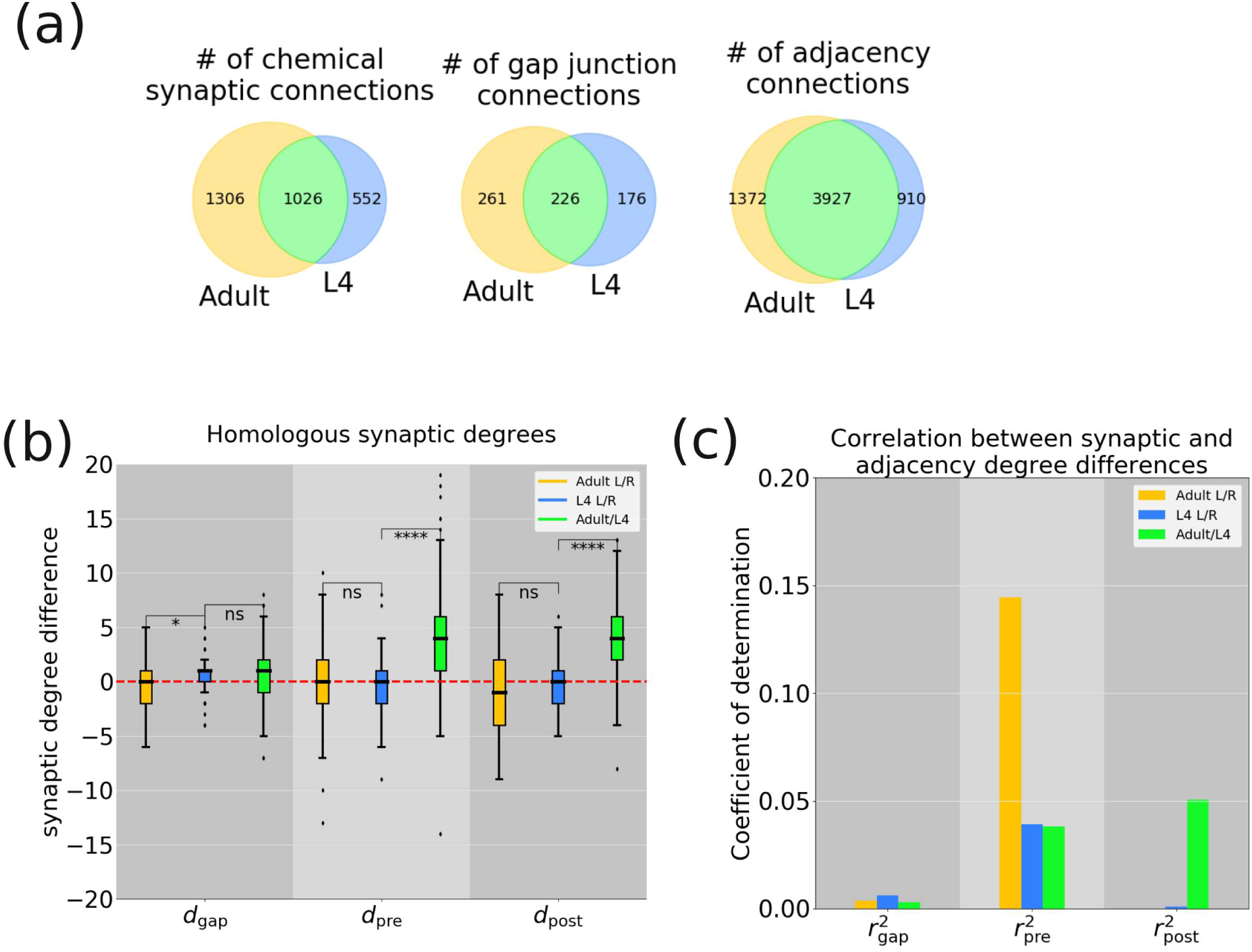
Variability in synaptic connectivity. (a) Venn diagrams showing the reproducibility of connections between the adult and L4 data sets for chemical synapses (left) gap junctions (center) and adjacency (right) connections.(b) Distribution of synaptic degree differences for homologous neurons. Compared gap junction (*d*_gap_), presynaptic (*d*_pre_) and postsynaptic (*d*_pre_) degrees. (ns) not statistically significant. (***) *p* < 0.05, *t*-test. (c) Correlation between synaptic and adjacency degree differences between homologous neurons. The coefficient of determination if given for gap junction (*r*_gap_), presynaptic (*r*_pre_) and postsynaptic (*r*_post_) correlations.

**Figure S4.**
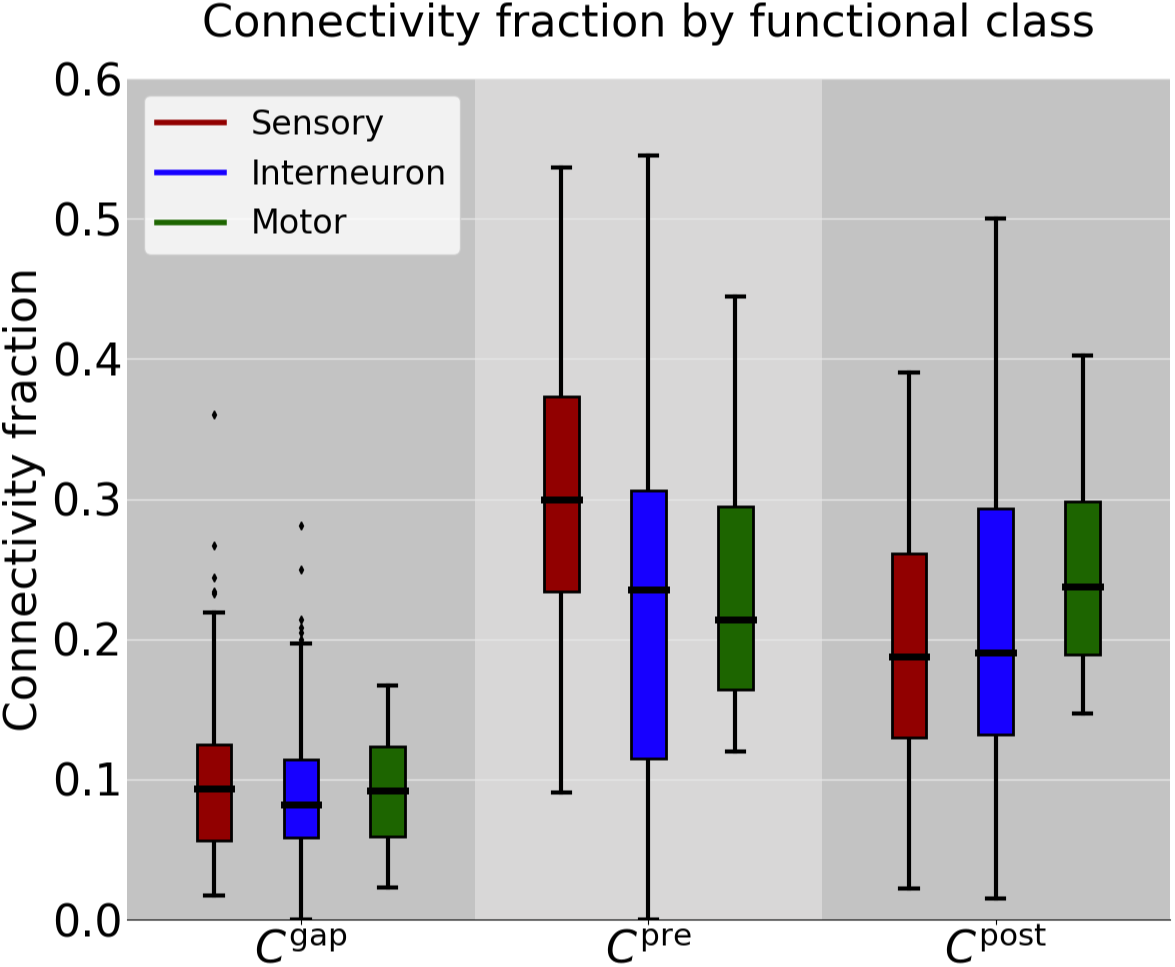
Connectivity fractions corrected for muscles. A significant fraction of motor neuron output and some sensory/interneuron output is onto muscle which is not included in our segmentation. To correct for this, we artificially added muscles to the adjacency list. If neuron *A* synapses onto muscle *M*, then the adjacency (*A*, *M*) was added to the adjacency list. Hence, every adjacency added has a corresponding synaptic connection. Therefore, the corrected connectivity fractions are likely overestimates because there are adjacencies with muscles that do not result in synapses.

**Figure S5.**
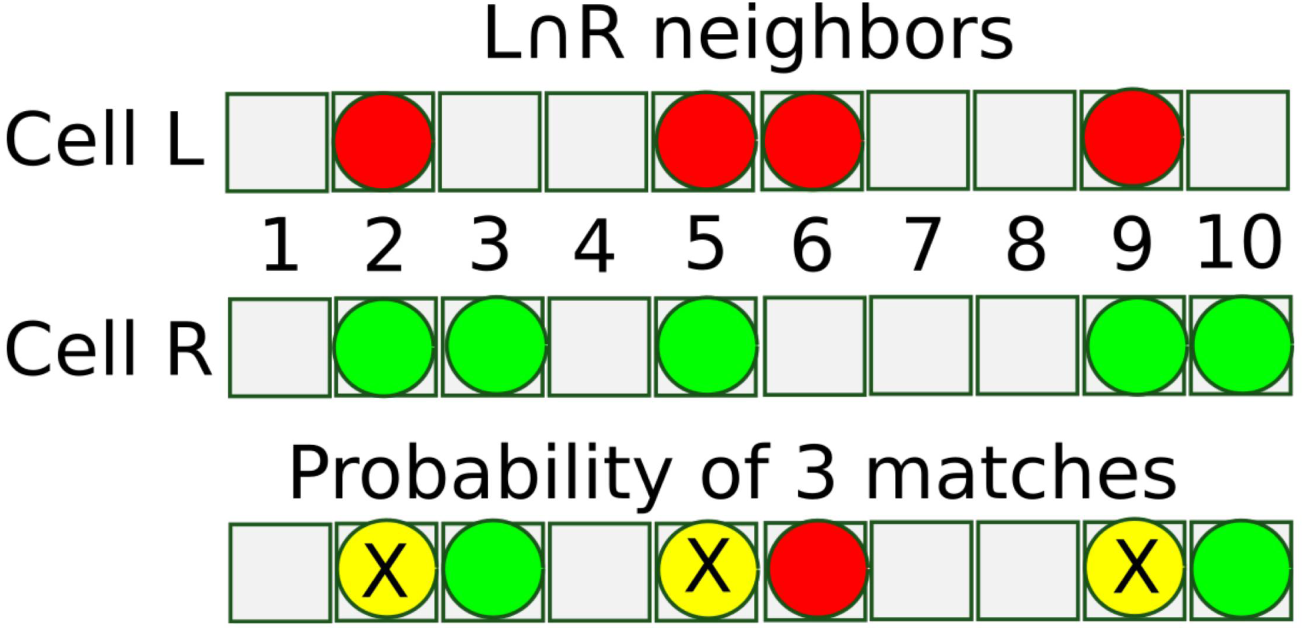
Specificity probability. Illustration of how specificity probability is computed. Suppose cell L and cell R have 10 common neighbors (numbered boxes); cell L makes 4 synaptic connections (red circles) onto neurons in the common neighborhood; and cell R makes 5 synaptic connections (green circles) onto neurons in this set. If cells L and R have 3 common synaptic connections (X on yellow circles in the bottom row), cells L and R make 1 and 2 additional (asymmetric) synaptic connections, respectively. The numerator of the specificity probability (Equation (6)) counts the number of possible combinations of non-overlapping yellow, red and green circles (here denoting symmetric L and R, L asymmetric and R asymmetric connections) among a set of (here 10) targets. The denominator counts the number of combinations of 4 in 10 red circles (for cell L) and the 5 in 10 green circles (for cell R) given the respective connectivity fractions.

**Figure S6.**
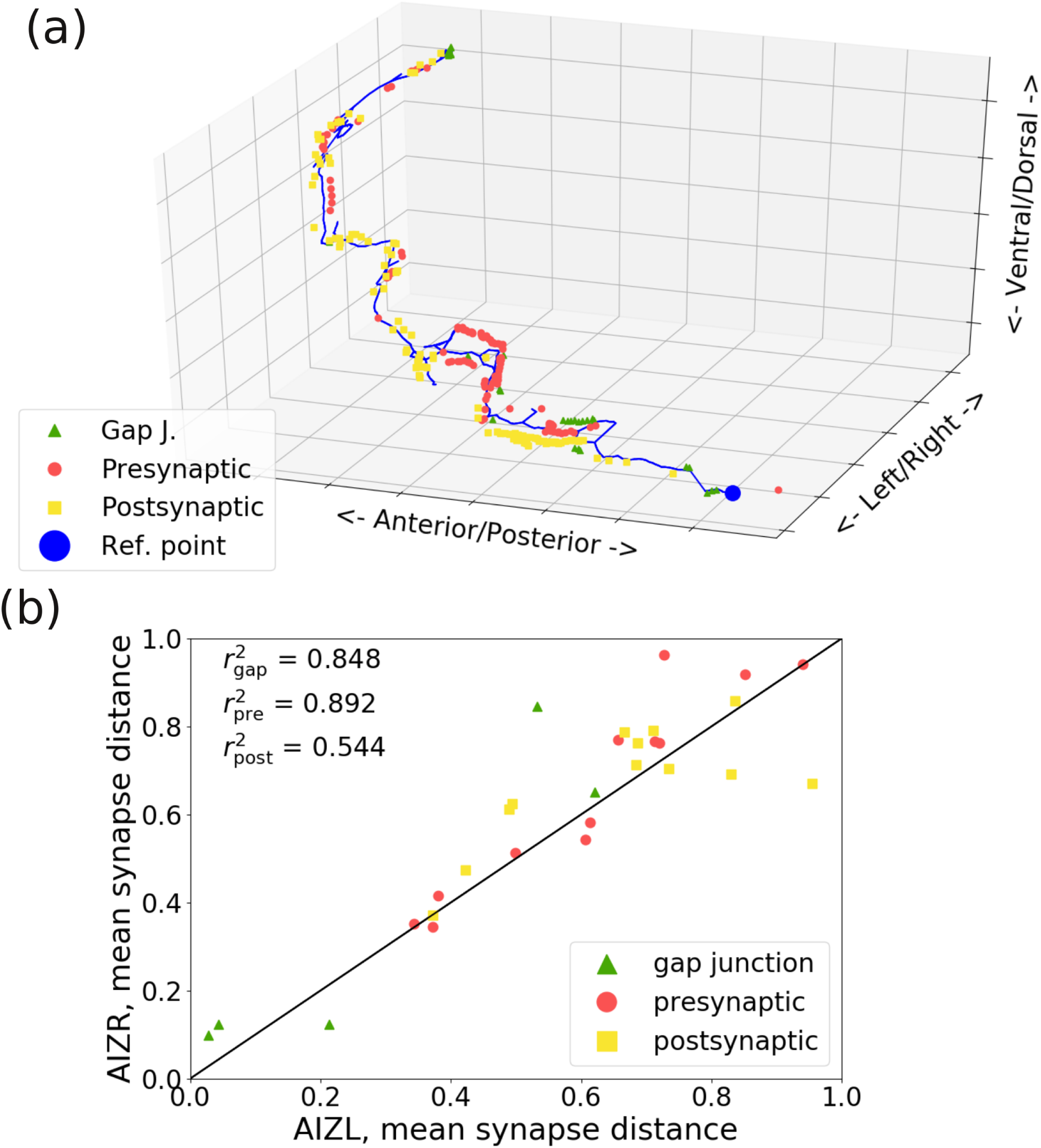
Correlation between mean synapse positions. (a) An example neuron skeleton using mesh contraction. Points of synaptic contact are plotted along the skeleton: gap junctions (green triangles), presynaptic (red circles) and postsynaptic (yellow squares). The reference point (large blue circle) is the point on the skeleton closest to the cell body. (b) Correlation of mean synapse position for homologous gap junctions, presynaptic and postsynaptic contacts for neurons AIZL and AIZR in the adult. Gap junction 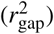, presynaptic 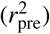 and postsynaptic 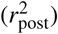 correlation coefficients.

**Figure S7.**
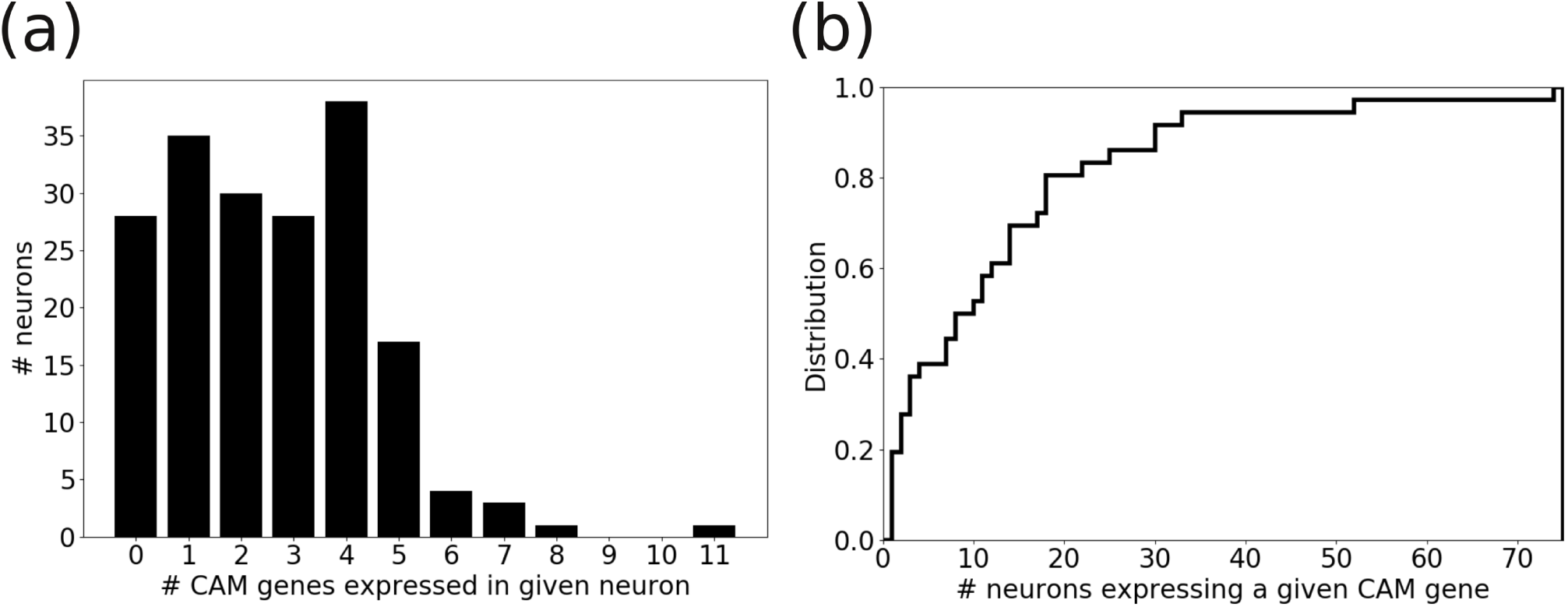
CAM expression in the nerve ring. (a) Histogram of the number of CAM genes expressed in an individual nerve ring neuron. (b) Cumulative distribution of the number of nerve ring neurons that express a given CAM gene.

**Figure S8.**
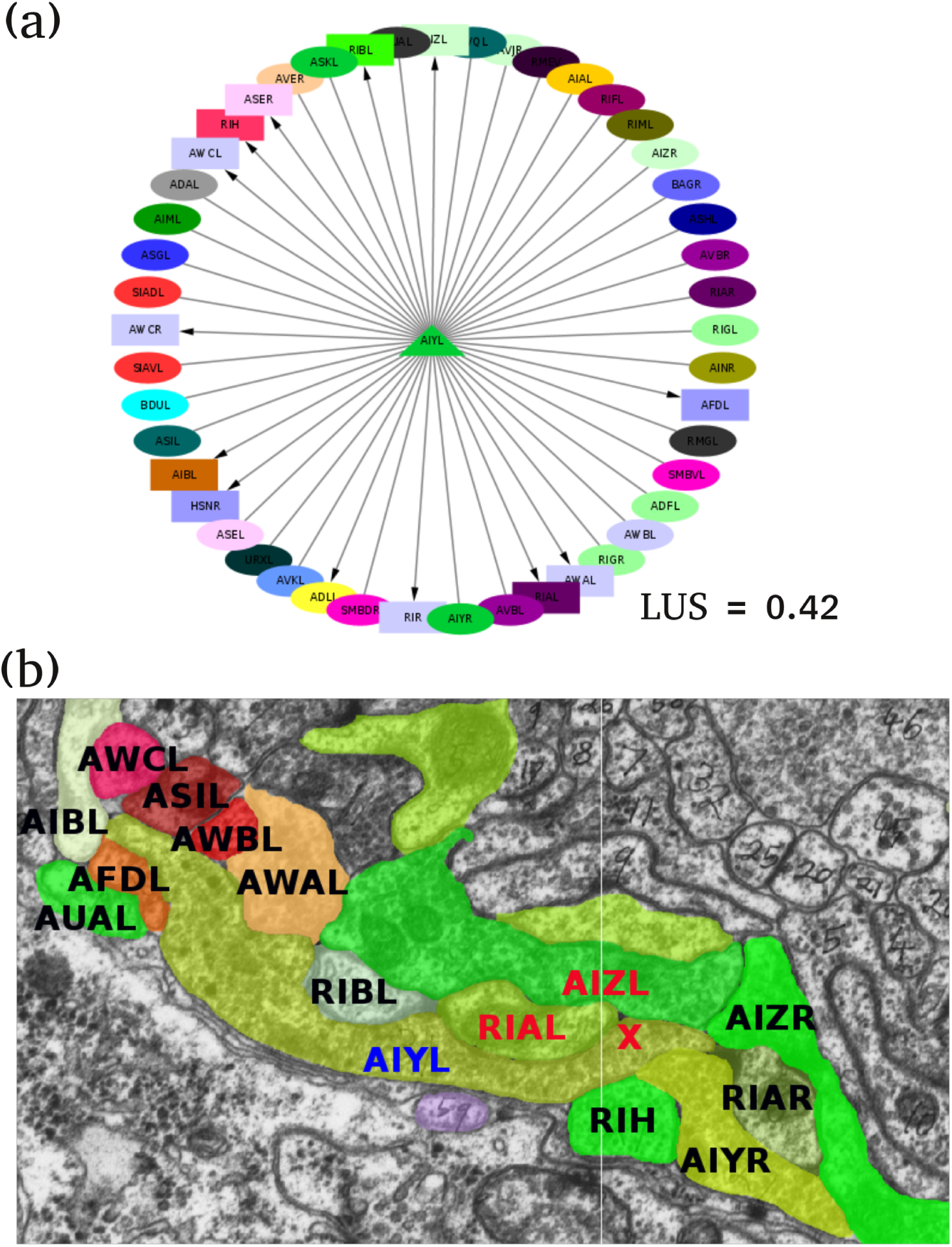
LUS calculation. (a) LUS for the WBE model. LUS for neuron AIYL (triangle) is illustrated. Postsynaptic partners (squares) and the remaining physically adjacent neurons (ellipses) are shown. Colors indicate the CAM expression label of the neuron. LUS is computed as 1 minus the fraction of squares whose color matches at least one ellipse. The LUS for AIYL is 0.42. LUS is computed similarly for the IE model. But in the IE model alternative splicing is simulated, therefore there is more diversity in CAM expression labels, i.e. neurons colors. (b) LUS for the SBE model. Rather than comparing expression labels at the cellular level, expression labels are compared at the subcellular level at synapse points. Shown is an EM section where AIYL (blue) synapses onto AIZL and RIAL (both labeled red). Synapse at the red ‘X’. The remaining physically adjacent neurons are shown labeled in black. For this particular synapse, the expression labels of AIZL and RIAL match the expression labels of AIZR and RIAR, respectively. Therefore, this synapse would decrease the LUS score of AIYL under the conditions of SBE model. Also worth noting is that this synapse is a particularly long synapse, occurring over 5 EM sections according to wormwiring.org.

**Figure S9.**
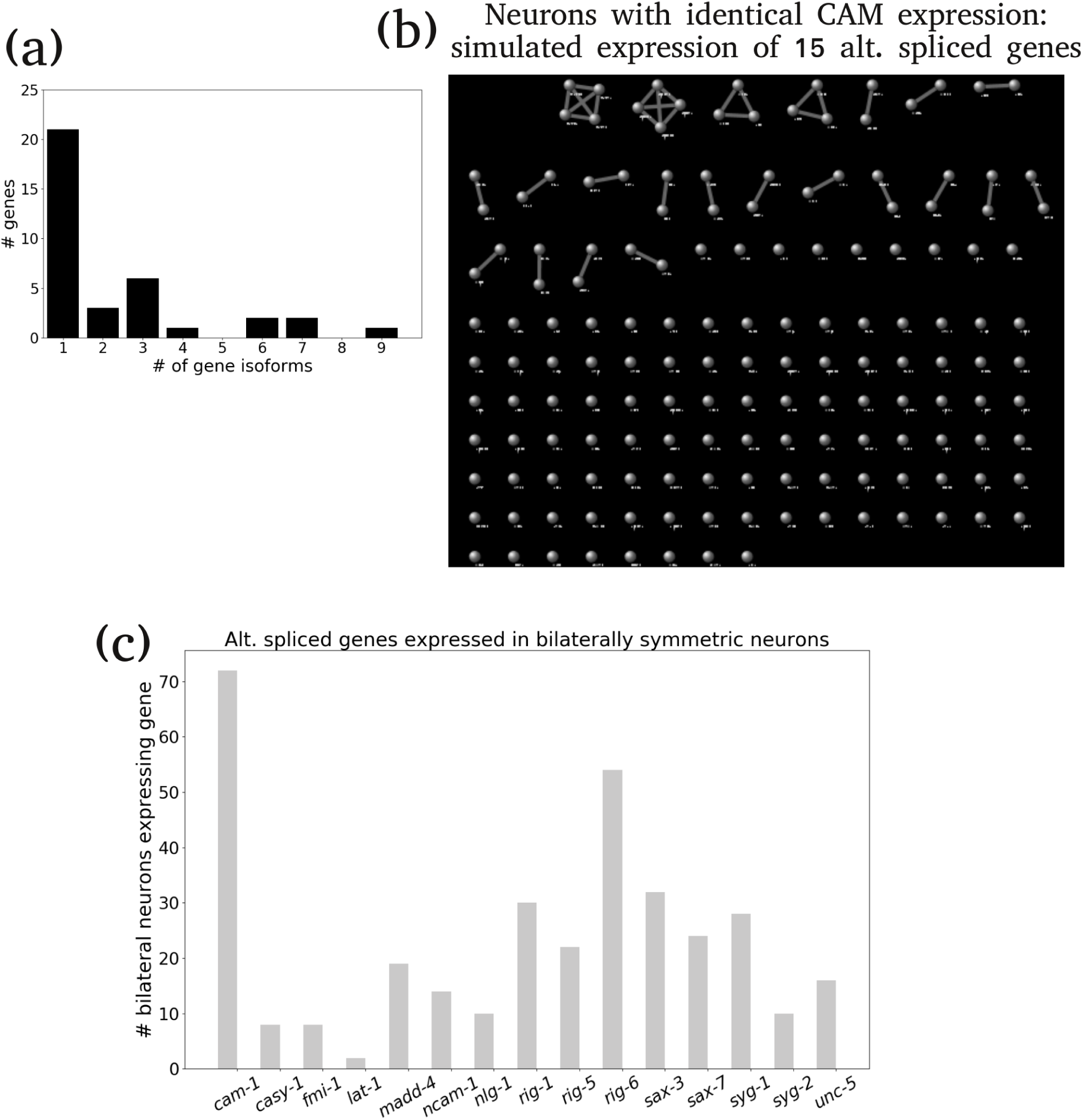
Alternative spliced CAM genes. (a) Histogram of the number of isoforms of CAM genes expressed in the nerve ring. (b) Graphs showing which neurons have the same CAM expression with alternative splicing. Neurons that have the same CAM expression pattern are connected by an edge. Neurons that have unique CAM expression (with simulated alternative splicing) are isolated nodes without an edge. Alternative splicing gives fewer clusters and thus more uniquely labeled neurons. (c) The number of neurons expressing alternatively spliced genes.

**Figure S10.**
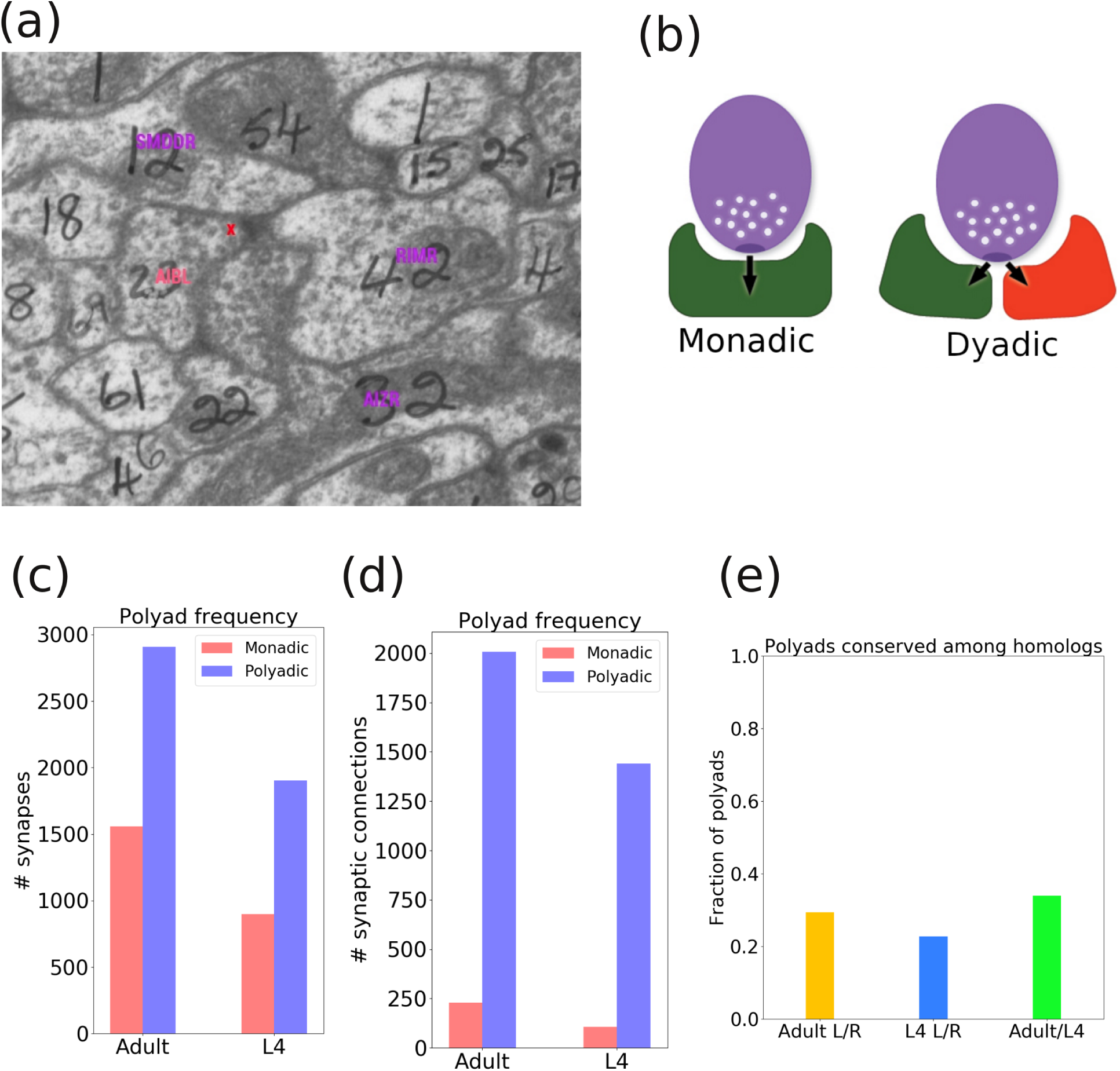
Polyadic connectivity. (a) Polyadic synapse from the adult data set. The synapse (marked by red x) has one presynaptic cell (AIBL) and three postsynaptic cells (SMDDR,RIMR and AIZR). (b) Cartoon illustrating the difference between a monadic and dyadic synapse. A monadic synapse has one postsynaptic cell, the dyadic synapse has two postsynaptic cells. Image taken from http://wormatlas.org. (c) Fraction of synapses that are polyadic. (d) Fraction of synaptic contacts that have at least one polyadic synapse. (e) Fraction of polyadic synapses that are conserved between homologous cells.

